# Cooperation, privatization and cheating in microbial exoenzyme synthesis: theoretical analysis in view of biotechnological applications

**DOI:** 10.64898/2026.05.29.728713

**Authors:** Tanvir Hassan, Shalu Dwivedi, Stefan Schuster, Anna Matuszyńska

## Abstract

This study presents a mathematical framework for investigating the dynamics of coexistence and competition among heterotrophic microbes across different time scales. Focusing on metabolic interactions, we examine how three strategies: public metabolizing, private metabolizing, and cheating, shape population behaviour. The framework integrates generalized Lotka–Volterra dynamics with evolutionary game theory to capture the effects of resource exchange, particularly glucose made available by public metabolizers and sucrose as a shared substrate driving population growth. Game-theoretic payoffs encode ecological costs and benefits, enabling analysis of frequency-dependent interactions among strategies. To capture evolutionary realism, we implement laboratory-inspired simulations in which strategies can switch between generations, mimicking mutation or phenotypic plasticity in microbial populations. These eco-evolutionary dynamics reveal conditions under which all three strategies coexist at interior equilibria and show how variation in growth advantages and, illustratively, phenotype-switching perturbations produce evolutionary shifts. Numerical analysis identifies ecological thresholds and fitness asymmetries that determine system robustness, long-term coexistence, and the persistence of a synthetic, cross-kingdom system linked by nutrient exchange. Together, these insights provide general principles for microbial coexistence and offer design guidelines for ecosystem engineering, biotechnological applications, and the construction of stable synthetic communities under ecological and evolutionary constraints.

## 1 Introduction

Microorganisms are omnipresent in nature, where they form diverse communities composed of genetically different strains or even a variety of species that interact and co-exist. However, they are not merely passive inhabitants: microorganisms constitute the functional backbone of most ecosystems. They regulate global biogeochemical cycles, drive primary production, and control nutrient turnover across environments ranging from the human colon microbiota [1] to cloud water [2] and soil consortia [3, 4]. Yet, how such diverse communities achieve stable and functional coexistence remains unclear. In parallel, there is growing interest in the rational design of synthetic microbial communities with defined composition and controlled interactions [5], for applications ranging from gut microbiome therapies to bioproduction and bioremediation [6, 7, 8]. This raises a key question: can such communities be designed to exhibit stable and predictable function?

To address it, we focus on microbial interactions. A key feature of microbial communities, both natural and synthetic, is their capacity to release a wide range of compounds into their surroundings. Microbes secrete metabolites, enzymes, and signalling molecules into their environment, thereby shaping the ecological landscape [9, 10, 11, 12]. These secretions often serve as the basis for cooperative, competitive, or neutral interactions [10]. For example, in mutualistic cross-feeding, the fungus *Laccaria bicolor* secretes trehalose to attract the bacterium *Pseudomonas aeruginosa*, which in turn produces thiamine to support fungal growth [13]. Some strains of *Saccharomyces cerevisiae* (baker’s yeast) secrete invertase into the periplasmic space or environment, allowing sucrose to be hydrolyzed outside the cell [14, 15]. This sugar then becomes accessible to both the producing and neighbouring non-producing cells.

To formalize these interactions, an economic perspective is often adopted, where microbes exchange resources analogously to agents in a market [16]. In this context, the production and sharing of extracellular compounds can be interpreted as public goods, giving rise to cooperation and exploitation dynamics that align naturally with evolutionary game theory (EGT) [17, 18, 19, 15, 20, 21, 22]. These dynamics have been extensively studied in evolutionary public goods frameworks [23, 24, 25]. In the classical cooperator–defector framework, individuals that produce shared resources are termed cooperators, while those that exploit them without contributing are defectors [18, 26, 27]. Optional Public Goods Games introduce a third strategy (loners) that avoid participation, enabling coexistence through cyclic or stable equilibria [28, 23, 29]. Ecological public goods models further incorporate density-dependent feedbacks, demonstrating that resource limitation and population dynamics can fundamentally alter cooperation outcomes [30, 27]. In parallel, it has been shown that cooperators and defectors with respect to exoenzyme production can coexist in a Snowdrift game (also known as Hawk-dove game) [14, 15]. More generally, eco-evolutionary analyses reveal that coupling strategy dynamics with population growth can give rise to oscillations, bifurcations, and multiple stable states [31]. However, these frameworks typically treat strategies abstractly and do not explicitly account for metabolite-mediated interactions or provide a direct mapping between ecological interaction coefficients and evolutionary payoffs.

Here, we adopt a metabolically grounded terminology and refer to cooperators as public metabolizers (*P*), and defectors as cheaters (*C*), following prior work emphasizing mechanistic descriptions of metabolite-mediated interactions [32]. Public metabolizers release extracellular enzymes or metabolites that benefit the community [33], and can be vulnerable to exploitation by cheaters, which avoid the energetic cost of enzyme production, but still reap the benefits of the communal pool of resources [34, 35]. Such exploitation can destabilize cooperative systems by reducing the net benefit of public goods production, potentially leading to population collapse or oscillatory dynamics [35, 34, 36, 37, 31]. A third strategy is given by private metabolizers (*M*), which internalize resource processing and avoid sharing [32, 34]. While they do not contribute to the communal pool and thus do not support cheaters, they still incur metabolic costs, distinguishing them from pure exploiters.

Ordinary differential equations (ODEs) based foundational approaches such as the Lotka–Volterra predator–prey models [38, 39] offer powerful frameworks to explore these dynamics systematically across ecological and microbial systems [40, 41]. While ODE-based ecological models capture density-dependent population dynamics, they do not directly encode strategy selection, although some game theoretical models have been based on ODEs [42, 43]. Conversely, EGT captures frequency-dependent selection but typically neglects explicit ecological feedbacks [44, 22]. This separation limits the ability to predict long-term outcomes in systems where metabolic interactions simultaneously determine both population growth and evolutionary fitness. In particular, Lindsay *et al*. [32] demonstrated that private metabolizers can invade and collapse systems of cooperators and cheaters under resource-explicit kinetics. However, this analysis primarily focused on invasion dynamics and does not provide a general characterization of coexistence or evolutionary stability across the full strategy space.

In this study, we develop a tripartite computational model to explore the ecological and evolutionary dynamics among three microbial strategies: public metabolizers, private metabolizers, and cheaters. The model builds on a generalized Lotka–Volterra framework [45], where population growth is influenced by access to resources, specifically glucose and sucrose (captured in Figure 1). This abstraction provides an explicit mechanistic mapping between density-dependent ecological interactions and evolutionary-game-theoretic payoffs, allowing ecological interactions to be interpreted as evolutionary strategies within a unified framework. This choice enables (i) direct mapping of per-capita growth rates onto evolutionary-game-theory payoffs, (ii) derivation of closed-form equilibrium expressions for all three strategies, (iii) systematic exploration of coexistence regimes via phase-flow diagrams on the strategy simplex, and (iv) a transparent protocol for estimating every model parameter from monoculture and pairwise co-culture experiments. Our main objective is therefore to determine the precise conditions under which all three strategies can stably coexist rather than collapse, with new mechanistic and evolutionary insights directly applicable to the rational design of synthetic microbial consortia. To incorporate the evolutionary dimension, we extended the model using EGT, allowing strategy frequencies to change based on relative fitness. This builds on prior game-theoretic analyses of microbial cooperation [14, 15], while explicitly incorporating metabolic privatization. It reveals not only which populations persist, but also how strategy composition changes under the coupled ecological–evolutionary dynamics, providing direct insight into the design of robust synthetic consortia.

**Figure 1:**
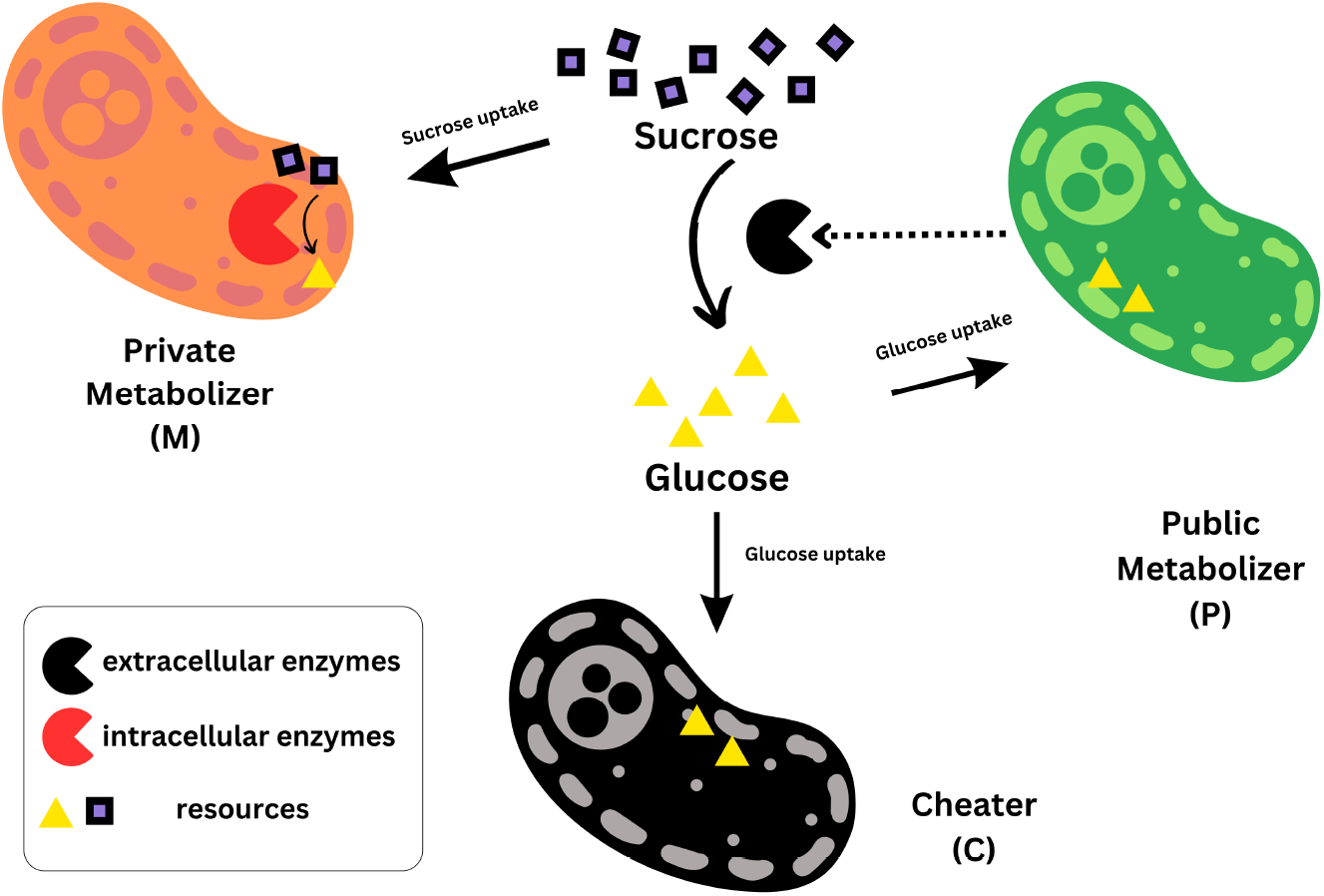
Schematic representation of a tripartite microbial community formalized based on their sugar-mediated interactions. All organisms are heterotrophic and utilize glucose (yellow triangle) derived from the breakdown of sucrose (black square). Public metabolizers (*P*) secrete extra-cellular enzymes (black Pac-Man symbol) to hydrolyse sucrose into glucose, contributing to a public pool of accessible resources. Cheaters (*C*) do not produce enzymes but exploit the publicly available glucose without incurring production costs. Private metabolizers (*M*) import sucrose for intracellular digestion using internal enzymes (red Pac-Man symbol), incurring metabolic costs while avoiding resource sharing. This scheme forms the conceptual basis for a mathematical model exploring the ecological dynamics of cooperation, exploitation, and metabolic privatization in microbial communities.

## 2 Modelling framework

Through this work, we model a closed, well-mixed batch culture, where three microbial strategies can be adopted: a public metabolizer (*P*), a cheater (*C*), and a private metabolizer (*M*) (Figure 1). Sucrose is continuously supplied as the sole carbon source. The public metabolizer secretes extracellular invertase, which hydrolyzes sucrose into glucose in the surrounding medium. This glucose is partially assimilated for its growth, while the rest becomes part of a common resource pool accessible to other community members.

### 2.1 Dynamic model of the tripartite population

To describe the ecological dynamics of these three strategies, we employ a generalized Lotka–Volterra framework that captures intrinsic growth, pairwise interactions, and density-dependent effects. Importantly, we assume that public metabolizers constitutively express invertase at a constant rate throughout their lifetime. Consequently, we do not explicitly model the dynamics of enzyme regulation, as done in other approaches [34, 37, 46], reducing the number of free parameters. Moreover, to make the point of cheating very clearly, we do not consider the possibility that cheaters could have a basal growth even when living alone. At the same time, it preserves ecological realism, since cheater growth remains directly dependent on the activity and density of public metabolizers.

The resulting system is governed by a set of coupled ODEs:

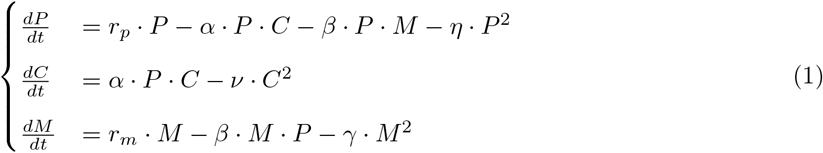

with seven effective parameters: intrinsic growth rates (*r*_*p*_, *r*_*m*_), an exploitation coefficient (*α*), an inter-strategy competition coefficient (*β*), and strategy-specific self-limitation coefficients (*η,ν, γ*). The interaction term *αPC* captures the growth benefit obtained by cheaters from public metabolizers and the corresponding loss imposed on the public metabolizer population. Thus, *α* quantifies exploitation intensity rather than cooperative investment per se. The quadratic self-limitation terms (−*ηP* ^2^, −*νC*^2^, −*γM* ^2^) arise as the effective density-dependent regulation in a closed, resource-limited batch culture. In such systems, net growth is ultimately constrained by the depletion of the sole carbon source (sucrose), the accumulation of metabolic by-products, and spatial interference, all of which scale nonlinearly with total biomass.

Rather than introducing an explicit dynamic equation for the shared resource pool, which would require additional parameters (uptake rates, half-saturation constants, enzyme turnover) that are difficult to identify from population time series alone, we adopt the standard reduced-form generalised Lotka-Volterra approximation widely used in microbial community modelling [45]. Critically, the three strategies incur distinct metabolic overheads: constitutive extracellular invertase secretion (P), intracellular enzyme synthesis and sucrose import (M), and zero production cost (C). Consequently, each experiences its own effective carrying capacity (*K*_*P*_ = *r*_*p*_*/η, K*_*M*_ = *r*_*m*_*/γ*), while cheater growth remains strictly exploitation-dependent. Allowing distinct coefficients for each strategy reflects differences in metabolic constraints and ecological niches among public metabolizers, cheaters, and private metabolizers. The cheater growth term contains no intrinsic rate *r*_*c*_ because, in the sucrose/invertase system modelled here, invertase-deficient mutants cannot hydrolyse sucrose and thus exhibit no net growth in monoculture [14, 47]. Their proliferation is strictly dependent on glucose released by public metabolizers, hence the growth rate is jointly determined by the exploitation coefficient *α* and the density of public metabolizers. However, to prevent uncontrolled proliferation, cheater growth is also regulated by density-dependent constraints*ν*.

Accordingly, the present model should be interpreted as a coarse-grained, phenomenological approximation rather than a fully resource-explicit mechanistic description. Within this framework, the parameters provide effective interaction strengths that can be estimated from experimental data, enabling tractable analysis of coexistence, stability, and evolutionary dynamics. This model is designed to be readily parameterizable from empirical data. To ensure broad applicability and experimental integration, we present a stepwise protocol for parameter estimation based on monoculture and pairwise co-culture experiments. This framework enables researchers to directly map ecological interactions onto measurable population dynamics.

#### 2.1.1 Model parameterization

Intrinsic growth rates of *P* (*r*_*p*_) and *M* (*r*_*m*_) can be estimated from exponential growth phase data of monoculture experiments. As population densities increase, density-dependent constraints (*η* and *γ*) capture the effect of crowding and resource limitation near the carrying capacity (Figure S2). These parameters are obtained by fitting the following simplified equations:

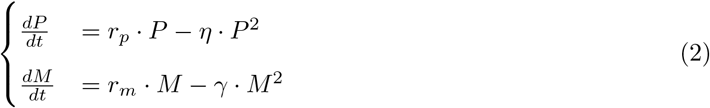

To further parameterize the interaction terms, we can analyze data from a pairwise co-culturing experiment, allowing us to isolate and quantify key interaction coefficients that define inter-strategy dynamics. The exploitation coefficient *α* is estimated by comparing the growth of the *P* in monoculture and in co-culture with *C* (Eq.3):

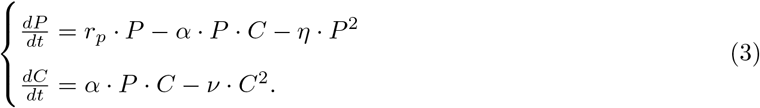

Similarly, the competition coefficient *β* quantifies interference between public and private metabolizers, both of which utilize the same external sucrose supply (Figure S2). Although they employ different processing strategies, this shared resource creates indirect competition. To estimate *β*, we compare the growth of *M* in monoculture to its dynamics in co-culture with *P* (Eq.4):

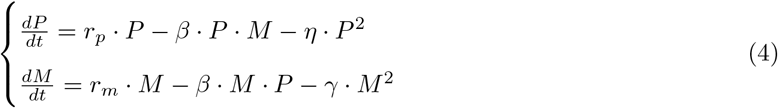

The three systems 2-4 provide minimalistic, yet interpretable, representations of microbial ecological dynamics, with clearly defined interaction terms and parameters grounded in empirical observables. It also lays the foundation for subsequent stability analyses, allowing us to explore co-existence regimes, dominance shifts, and resilience under varying environmental conditions.

The default parameter values used throughout the main-text simulations and figures are listed in Table 1. These values were chosen to produce biologically plausible growth rates and interaction strengths consistent with the sucrose/invertase system and are the values from which all phase-flow diagrams, sensitivity analyses, and evolutionary simulations in the main text are generated.

**Table 1:** Default parameter values used in the main simulations. Starting subpopulation sizes are set to 100 each.

| Parameter | Symbol | Default value |
| --- | --- | --- |
| Public metabolizer growth rate | $r_p$ | 0.5 |
| Private metabolizer growth rate | $r_m$ | 0.2 |
| Exploitation coefficient ($P \rightarrow C$) | $\alpha$ | 0.0002 |
| Competition coefficient ($P \leftrightarrow M$) | $\beta$ | 0.0001 |
| Density-dependent self-limitation (public) | $\eta$ | 0.0001 |
| Density-dependent self-limitation (cheater) | $\nu$ | 0.0001 |
| Density-dependent self-limitation (private) | $\gamma$ | 0.0001 |

#### 2.1.2 Numerical implementation and interactive model exploration

All simulations were implemented in Python and are available in the accompanying GitHub repository: https://github.com/Computational-Biology-Aachen/FightClub/. In addition to the reproducible notebooks, we provide a browser-based implementation of the tripartite model at: https://computational-biology-aachen.github.io/mxl-web/tripartite. This interactive version allows users to vary model parameters and initial conditions directly in the browser and to inspect the resulting population trajectories without local software installation. The web interface is intended to support transparent model exploration, parameter sensitivity inspection, and communication of model assumptions to experimental collaborators, exploring the web-native modelling approach described in [48].

### 2.2 Evolutionary Game Theory and Microbial Strategies

The outcome of interactions of subpopulations exhibiting different metabolic strategies is shaped by natural selection and often depends on the frequency of competing strategies, which EGT captures effectively. Our study considers a three-strategy game, where each of the three mechanisms represents a distinct subpopulation. To assign payoffs appropriately, our model accounts for both the benefits and drawbacks experienced by each subgroup.

#### 2.2.1 Three-strategy evolutionary game

The payoff illustrates the complex relationships and fitness outcomes among these strategies within a microbiological community. From an evolutionary perspective, natural selection often favours cheaters, as they benefit from shared resources without contributing to their production. This typically leads to a decline in the *P* population. The success of each strategy is thus dependent on population densities and the payoff of that strategy [49].

In continuous-time evolutionary game theory, the payoff of each strategy is defined as its per-capita growth rate, i.e., 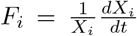 for strategy *i* with abundance *X*_*i*_. Consequently, the payoff functions given above are obtained directly by dividing the right-hand sides of the Lotka–Volterra system (1) by the respective population densities. This equivalence guarantees that any interior ecological fixed point (where 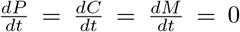) automatically satisfies *F*_*P*_ = *F*_*C*_ = *F*_*M*_ = 0, which is the equilibrium condition for an interior fixed point in continuous-time evolutionary game theory. We note that using per-capita growth rates as payoffs is standard in ODE-based evolutionary game models [43], and that all payoffs vanish at the ecological equilibrium, a direct consequence of steady-state growth rates equalling zero rather than an indication of equal but non-zero payoff values Appendix C. Therefore, the player payoffs for different strategies can be expressed as:

i. Public Metabolizers:

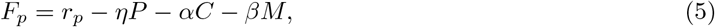
ii. Cheaters :

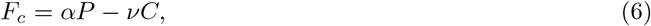
iii. Private Metabolizers :

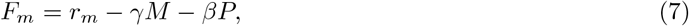

In this model, payoff functions are directly mapped to the intrinsic growth rates of individual strategies, serving as quantitative measures of fitness that govern population dynamics. A strategy with a higher payoff increases in relative abundance over time, shifting community composition toward public–cheater or private dominance depending on which growth rate advantage prevails.

## 3 Results

To investigate the ecological and evolutionary dynamics of our tripartite microbial system, we first performed a series of numerical simulations based on a generalized Lotka–Volterra framework. This approach enabled us to systematically vary key parameters, such as intrinsic growth rates and interaction strengths, and examine how these shape population trajectories over time. We also performed a sensitivity analysis that quantifies the impact of perturbing key model parameters on steady-state outcomes. This approach enables us to identify which interactions have the strongest influence on system behaviour and long-term community structure.

### 3.1 Sensitivity analysis

A systematic one-at-a-time sensitivity analysis was performed by varying each parameter individually while holding all others at their baseline values (Table 1). Each parameter was perturbed multiplicatively over the range 0.5–1.5× its nominal value (i.e. −50%, −20%, +20%, +50%). For each perturbation, the system was simulated from fixed initial conditions, and the resulting changes in population abundances were quantified relative to the baseline simulation. Sensitivities were computed either as relative changes in steady-state values *P* ^*^, *C*^*^, and *M*^*^ (when convergence was reached) or, where applicable, as relative changes in time-integrated population abundance (Supplementary Figure S4).

The resulting sensitivities are summarised in Supplementary Table S1. The system is most sensitive to the intrinsic growth rates *r*_*p*_ and *r*_*m*_. Increasing *r*_*p*_ strongly enhances the public–cheater subsystem, leading to a simultaneous increase in *P*^*^ and *C*^*^ and a reduction in *M*^*^. Conversely, increasing *r*_*m*_ favours private metabolizers at the expense of the public-cheater pair. The exploitation coefficient *α* and the private self-limitation coefficient *γ* also exert substantial influence: higher *α* promotes cheaters while suppressing private metabolizers, whereas higher *γ* reduces *M*^*^ and indirectly benefits *P* ^*^ and *C*^*^. The interaction coefficient *β* and the density-dependent terms *η* and *ν* have weaker but still non-negligible effects.

To assess robustness, results were evaluated across multiple perturbation magnitudes (±20% and ±50%), which yielded consistent qualitative trends (Supplementary Table S1). Together, these findings indicate that coexistence and regime structure are primarily governed by growth-rate asymmetries and exploitation strength, consistent with the analytically derived feasibility conditions (subsection 3.6) and the phase-flow analysis (Figure 4).

### 3.2 Impact of the growth rate

We started the analysis of our model by testing various growth rates while keeping other parameters constant, thereby allowing us to test the dynamics of species interactions within a shared environment. In scenarios involving public and private metabolizers, stable coexistence is critically dependent on the balance between their respective growth rates (*r*_*p*_ and *r*_*m*_). The public metabolizer plays a vital role in maintaining community function, potentially by producing shared resources or maintaining favourable environmental conditions that benefit the total community. However, when the growth rate of the private metabolizer becomes excessively high, it can dominate the system by consuming a disproportionate share of resources or suppressing the growth of the public metabolizer. This imbalance may result in system instability, and thereby leading to the extinction of both the public metabolizer and the cheater population (Figure 2).

**Figure 2:**
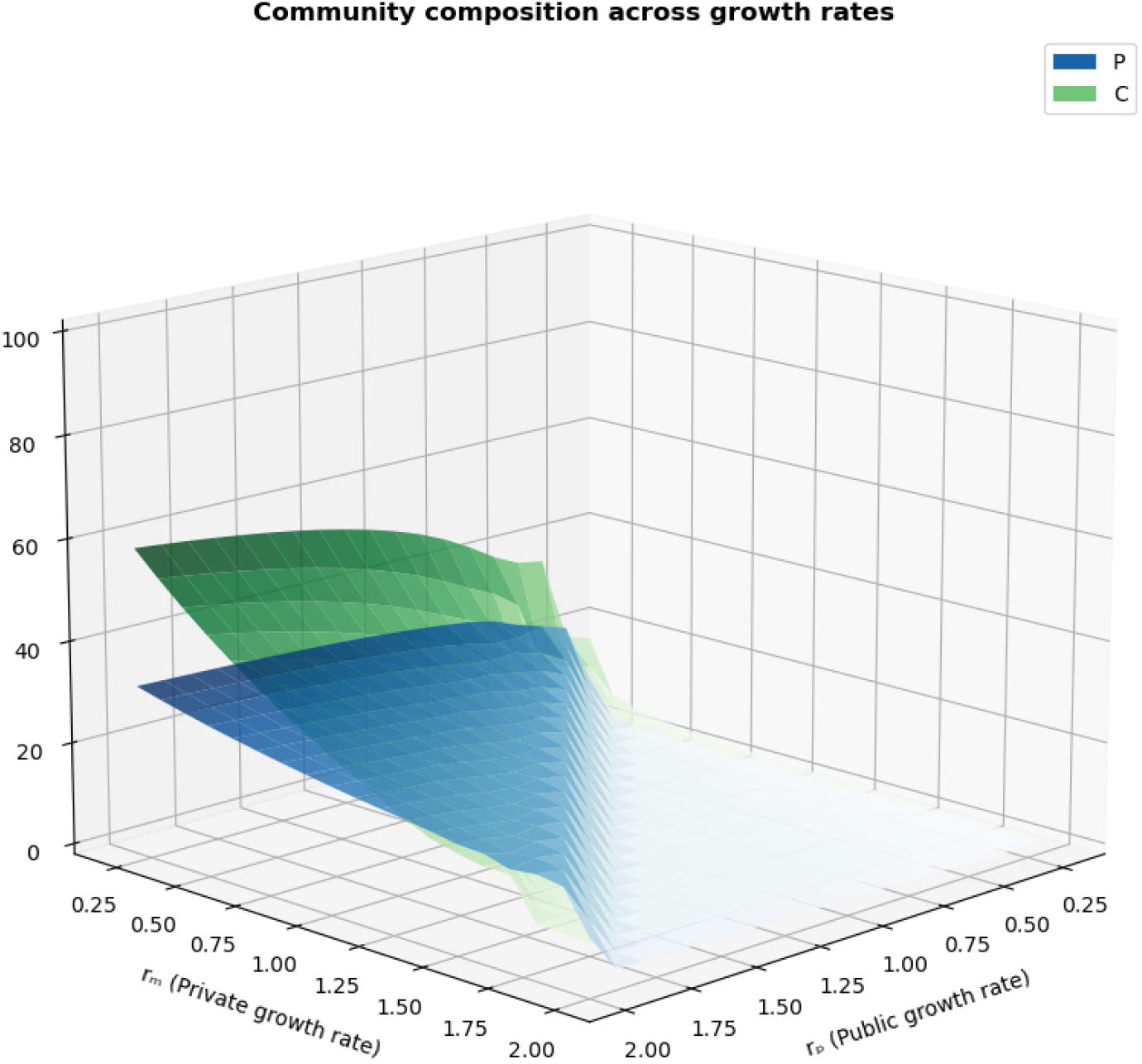
Impact of different growth conditions of *P* and *M* on population abundance. In this figure, we systematically investigate how changes in the intrinsic growth rate parameters *r*_*p*_ and *r*_*m*_ influence the ratio of *P* and *C* in the total population.

Numerical simulations demonstrate that stable coexistence among public, cheater, and private metabolizers is highly sensitive to the balance of their growth rates. In particular, the public metabolizer must sustain a growth rate sufficiently high to maintain its own population while also supporting the cheater population that relies on shared resources. However, this growth advantage must be carefully constrained: when the public metabolizer’s growth rate exceeds approximately 2 times that of the private metabolizer, competitive pressures lead to the decline or extinction of the private metabolizer (Figure 2). Conversely, if the public metabolizer’s growth rate falls below roughly 1.2 times that of the private metabolizer, neither the public nor the cheater populations can persist, resulting in the dominance of the private metabolizer. Therefore, a narrowly defined growth rate ratio, between 1.2 and 2 times higher for public metabolizers, is essential to ensure balanced and stable coexistence among all three metabolic strategies.

Further analysis of population dynamics reveals that excessive growth by any single population leads to increased density-dependent self-limitation, reducing net growth rates across the community. Similarly, in the absence of one population, the remaining species may initially increased growth, but subsequently encounter constraints arising from density-dependent effects encoded in the model. While these effects may reflect underlying limitations such as resource competition in real systems, the present model captures them implicitly through effective interaction terms rather than explicit resource dynamics. Collectively, these results demonstrate how variations in growth and interaction parameters can substantially influence the persistence and composition of microbial communities.

### 3.3 Impact of initial population size

The dynamic interplay among public metabolizers, cheaters, and private metabolizers in microbial communities reflects a finely balanced ecological system, significantly influenced by initial population densities in the short term, yet exhibiting long-term robustness under balanced growth conditions. During the early phases of community development, differences in initial population composition influence short-term dynamics. When cheaters are introduced at low initial densities, public metabolizers can sustain their growth while simultaneously supporting the cheater population through the provision of shared resources (Figure S1a). Conversely, elevated initial densities of cheaters accelerate resource depletion, prompting a temporary decline in public metabolizers and destabilizing the early stages of community structure (Figure S1b). Despite early fluctuations, our long-term simulations consistently demonstrate convergence toward a common steady-state equilibrium across all tested initial conditions, provided that growth parameters remain balanced. Specifically, after prolonged co-culture periods, initial differences diminish, resulting in nearly identical stable populations for all three strategies (Figure S1c). These findings suggest that, under balanced growth conditions, initial population distributions exert limited influence on long-term outcomes when growth parameters are optimally balanced, highlighting the inherent robustness of the tripartite coexistence to initial perturbations.

Notably, our simulations also reveal a transient ecological advantage for private metabolizers under conditions of elevated initial cheater densities. In scenarios where cheaters initially predominate, their rapid overexploitation of shared resources suppresses public metabolizers, thereby reducing competitive pressure and allowing private populations to expand more rapidly (Figure S1b). While this shift gives rise to short-term deviations from typical community dynamics, long-term equilibria consistently re-establish tripartite coexistence once growth parameters stabilize, highlighting the system’s resilience and adaptive capacity.

Collectively, these results highlight that while short-term community dynamics are sensitive to initial conditions and population imbalances, the long-term stability and coexistence of these microbial strategies are primarily determined by balanced growth conditions rather than initial community composition. Such insights are crucial for designing and managing stable synthetic microbial ecosystems capable of sustaining robust performance across a range of starting configurations and environmental contexts.

### 3.4 Impact of the interaction coefficients

The ecological interactions within microbial communities are significantly shaped by parameters *α* and *β*, which govern the benefit-sharing and competition dynamics in the system. This study explores how changes in *α* and *β* independently affect community dynamics, holding other parameters constant. The current analysis specifically considers a previously balanced community, characterized by a higher intrinsic growth rate of public metabolizers.

A higher *α* value indicates greater resource exploitation by cheaters from public metabolizers. When *α* exceeds *β*, cheater populations grow substantially, often at the expense of public metabolizers due to intense resource diversion. This imbalance can destabilize the community, significantly reducing the population of public metabolizers crucial for ecosystem stability. However, excessively high values of *α* can ultimately also reduce cheater populations, as their survival directly depends on the availability of public metabolizers. Parameter *β* quantifies competition between public and private metabolizers for shared nutrients. Higher *β* values indicate intense competition, severely constraining the growth of both populations. When *β* significantly exceeds *α*, the private metabolizer population cannot compete effectively and is driven towards extinction. Conversely, lower *β* values permit independent growth, but risk unchecked proliferation of private metabolizers, potentially destabilizing the community.

The Interactions between *α* and *β* critically define coexistence conditions among public, private, and cheater populations. From the analysis and visual inspection of the provided figures, the optimal scenario for stable coexistence is observed when *α* is moderately higher than *β*. Under these conditions, cheaters thrive without causing substantial harm to public metabolizers, and public and private populations maintain competitive yet balanced growth. Deviations from this optimal balance, particularly excessively high *β* or disproportionately high *α*, disrupt community stability, resulting in dominance or extinction events. Figure 3 illustrates these dynamics, emphasizing parameter ranges that best support stable coexistence among all three microbial populations.

**Figure 3:**
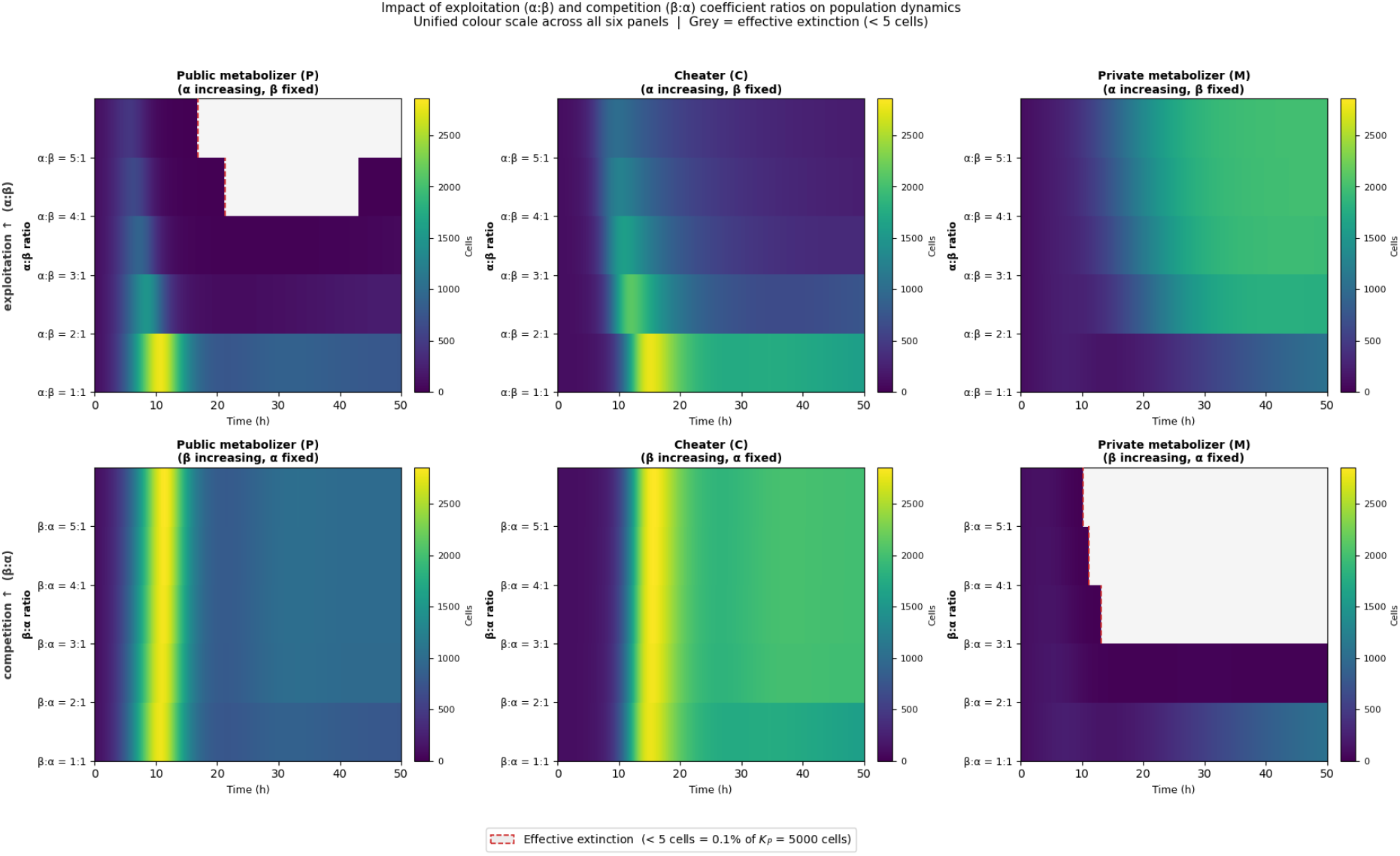
Impact of interaction coefficient ratios on population dynamics of the tripartite microbial community. Heatmaps show the temporal evolution of public metabolizer (P, left), cheater (C, centre), and private metabolizer (M, right) population densities under systematic variation of the exploitation (*α*) and competition (*β*) coefficients. Upper row: the exploitation coefficient *α* is scaled from 1x to 5× its baseline value (*α*:*β* ratios 1:1 to 5:1) while *β* is held fixed, reflecting increasing cheater benefit relative to competitive cost. Lower row: the competition coefficient *β* is scaled from 1x to 5x (*β*:*α* ratios 1:1 to 5:1) while *α* is held fixed, reflecting increasing resource competition between public and private metabolizers. A unified colour scale spanning 0–2856 cells is applied across all six panels to enable direct cross-panel comparison. Grey shading marks effective extinction, defined as population density falling below 0.1% of the public metabolizer carrying capacity (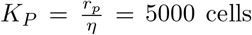; threshold = 5 cells); the dashed red line indicates the time of first crossing of this threshold. All simulations use the default parameter set (Table 1) with initial conditions P_0_ = C_0_ = M_0_ = 100 cells.

### 3.5 Stability of a Two-Strategy Coexistence

In populations with two co-existing strategies, the third strategy can often invade due to the interplay of exploitation and competition dynamics. For example,when public metabolizer and cheater coexist (*P, C >* 0, *M* = 0), the fitness of public is *F*_*p*_ = *r*_*p*_ −*αC* −*ηP*, and the fitness of cheater is *F*_*c*_ = *αP* −*νC*. Private metabolizer can invade this system if its fitness, *F*_*m*_ = *r*_*m*_ − *βP* − *γM*, exceeds both *F*_*p*_ and *F*_*c*_, which occurs when *r*_*m*_ *> r*_*p*_ − *αC* and *r*_*m*_ *> αP* −*νC*. In coexistence between public metabolizer and private metabolizer (*P, M >* 0, *C* = 0), the fitness of *P* is *F*_*p*_ = *r*_*p*_ − *ηP* − *βM*, and the fitness of *M* is *F*_*m*_ = *r*_*m*_ −*γM* −*βP*. Cheaters can invade this system by exploiting public metabolizers, achieving fitness *F*_*c*_ = *αP >* 0, provided *α > η*. Lastly, in a system with cheaters and private metabolizers (*C, M >* 0, *P* = 0), where *F*_*c*_ = *αP* −*νC* and *F*_*m*_ = *r*_*m*_ −*γM* −*βP*, public metabolizers can invade if *r*_*p*_ *> r*_*m*_ −*βM* +*γM*. These mathematical relationships reveal that the coexistence of two strategies is typically unstable, as the third strategy can always exploit competitive or cooperative dynamics to establish itself, indicating that stable coexistence, when it occurs, requires an interior equilibrium involving all three strategies.

### 3.6 Stability of the interior equilibrium

The dynamics work by increasing the proportion of strategies that yield higher payoffs. For instance, if the public strategy offers greater fitness compared to private, the population will evolve toward a public-cheater equilibrium. Similarly, if the private strategy offers a higher payoff than the public one, the system will shift toward a private equilibrium. Mutual cheating is generally discouraged, because it produces negative payoffs. In the case of public-cheater-private mixed equilibrium, the populations have a lower fitness because of a relative loss of cooperation and competition, which reduces the effectiveness of this strategy. Therefore, the system moves toward mixed stable equilibria.

The coexistence of all three strategies (*P, C, M >* 0) corresponds to an interior fixed point of the Lotka– Volterra system (Eq. 1), where 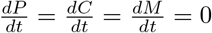. As shown in the subsection 2.2, this ecological fixed point is mathematically identical to the point at which all three payoffs are equal (*F*_*P*_ = *F*_*C*_ = *F*_*M*_ = 0), constituting a stable interior evolutionary equilibrium of the system. Solving these steady-state conditions yields the interior equilibrium densities derived in Appendix C:

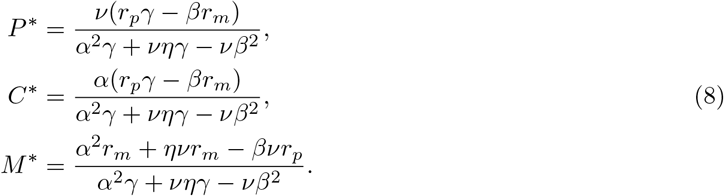

The local stability of this interior equilibrium is not implied by the existence of the fixed point itself. It is assessed separately by evaluating the Jacobian matrix of the population dynamics at (*P* ^*^, *C*^*^, *M*^*^), as shown in Appendix D. Across the biologically feasible parameter region in which all three equilibrium densities are positive, the Jacobian eigenvalues have negative real parts, confirming local asymptotic stability.

The feasibility limits of the interior equilibrium are most easily interpreted by comparison with the boundary equilibria of the system (explicitly in Appendix E). These equilibria correspond to ecological regimes in which one or more strategies are absent, including the *M*-only state, the *P*-only state, the *P*– *C* subsystem, and the *P*–*M* subsystem. Biologically, these boundary equilibria describe the alternative outcomes obtained when coexistence fails: collapse of the public-good system toward private metabolism, exclusion of private metabolizers by the public–cheater subsystem, or coexistence of public and private metabolizers in the absence of cheaters. The vertices and edges approached in the phase-flow diagrams in Figure 4 correspond to these boundary equilibria.

**Figure 4:**
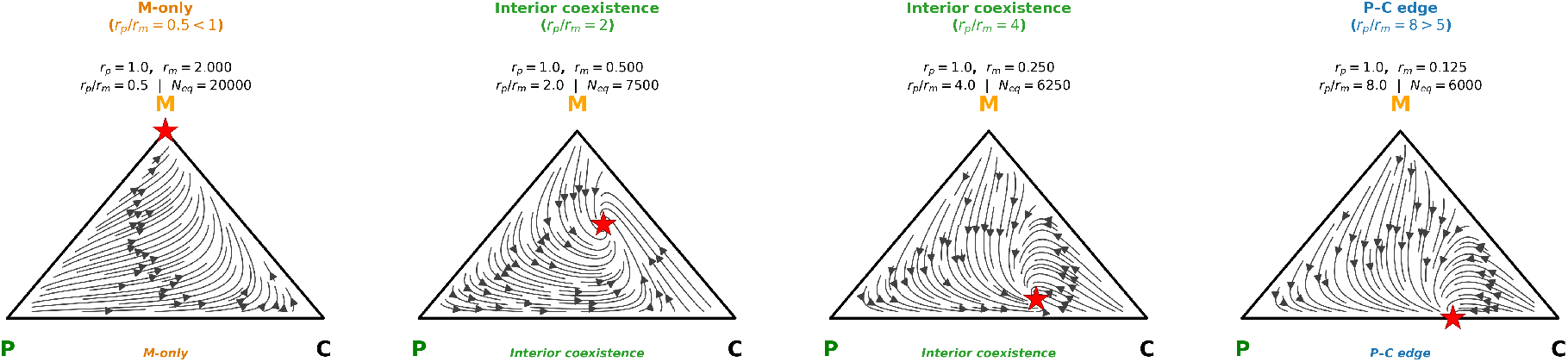
Projected phase portraits showing payoff-associated regime shifts in the system. Each triangle represents all possible relative abundances of three strategies (*P*–*C*–*M* simplex), with *N*_eq_ being the total abundance of the analytically derived attractor for the respective parameter set. With the default parameters, the feasibility window for interior coexistence corresponds to 1 *< r*_*p*_*/r*_*m*_ *<* 5. The four panels illustrate representative regimes as *r*_*p*_*/r*_*m*_ increases: convergence to the *M*-only boundary equilibrium (*r*_*p*_*/r*_*m*_ = 0.5), two feasible interior coexistence equilibria (*r*_*p*_*/r*_*m*_ = 2 and *r*_*p*_*/r*_*m*_ = 4), and convergence to the *P*–*C* boundary equilibrium (*r*_*p*_*/r*_*m*_ = 8). Convergence to an interior point indicates stable coexistence of all three strategies, with *P* ^*^ *>* 0, *C*^*^ *>* 0, and *M*^*^ *>* 0. Red stars 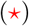 mark the analytically derived equilibrium composition for each parameter regime. Local stability of the interior equilibrium is assessed by Jacobian analysis in Appendix D.

These equilibrium expressions are biologically feasible only when all three quantities (*P* ^*^, *C*^*^, *M*^*^) are strictly positive, which requires the growth rates to satisfy:

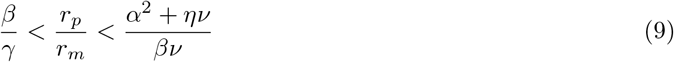

(derived in Appendix D). When *r*_*p*_/*r*_*m*_ falls below *β*/*γ, M*^*^ remains positive but *P* ^*^ and *C*^*^ become negative, meaning the interior equilibrium is not biologically realised and the system instead converges to the *M*-only boundary equilibrium. When *r*_*p*_/*r*_*m*_ exceeds 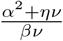, *M*^*^ becomes negative and the system converges to the *P* −*C* subsystem equilibrium. These transitions represent transcritical bifurcations: as *r*_*p*_/*r*_*m*_ crosses each threshold, the interior equilibrium collides with a boundary equilibrium and exchanges stability with it [41]. The local stability of each boundary equilibrium can be assessed by standard sub-Jacobian analysis; see, for example, Strogatz [41].

Substituting the interior equilibrium expressions (Equation 8) into the payoff functions (Eqs. 5–7) confirms that *F*_*P*_ = *F*_*C*_ = *F*_*M*_ = 0 (algebraic verification is provided in Appendix C). Local stability of this interior equilibrium was assessed via the Jacobian matrix of the system evaluated at (*P* ^*^, *C*^*^, *M*^*^) (Appendix D); numerical evaluation across the biologically relevant parameter ranges (Figure 4) shows that the eigenvalues have negative real parts whenever the equilibrium proportions are positive, confirming asymptotic stability.

Figure 4 illustrates these qualitative transitions across parameter space: each panel is a phase portrait for a specific (*r*_*p*_, *r*_*m*_) combination, and together they show how the attractor shifts from an interior coexistence point to a boundary equilibrium as the growth-rate ratio crosses the analytically derived thresholds in Appendix D. When one strategy has a strong growth advantage, the streamlines move toward the corresponding vertex, leading to complete dominance. In contrast, when payoffs are more balanced, the flow converges toward an interior point or along an edge, representing a stable coexistence of multiple strategies.

Increasing density-dependent parameters shifts the interior equilibrium toward the *P* −*M* edge, reflecting how ecological crowding suppresses cheaters persistence. From any such baseline, growth rate asymmetries determine the direction of further equilibrium movement: raising *r*_*p*_ relative to *r*_*m*_ drives the system toward the *P* −*C* edge, while increasing *r*_*m*_ drives it toward the *M* apex. Once *r*_*p*_ falls below the threshold *βr*_*m*_*/γ*, public metabolizers lose positive payoff, cheaters collapse with them, and the system converges to M-only dominance. These transitions correspond to the transcritical bifurcations at the boundaries of the feasibility window and are visible in the phase portraits of Figure 4.

### 3.7 Phenotype-switching perturbation simulation

Under the appropriate ecological conditions identified in the previous sections, the three strategies can coexist at locally stable interior equilibrium. While such equilibria may be associated with reduced average fitness due to the combined costs of cooperation and exploitation, they represent a dynamic balance in which no single strategy can dominate persistently. Here, the success of each strategy is determined not only by its intrinsic growth rate but also by its interactions with the others.

To further capture evolutionary realism, we employed laboratory-inspired evolutionary simulations that allowed individual strategies to change between generations, mimicking realistic microbial adaptation through mutation or phenotypic switching.

After each ecological growth phase, we imposed a fixed strategy-transition step in which a prescribed fraction *s*_*t*_ of the private-metabolizer population was reassigned to either the public-metabolizer or cheater strategy. For the private-to-public transition, the update was

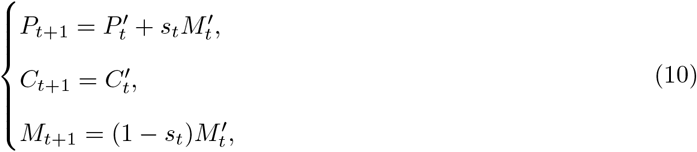

where 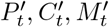 denote the population sizes after one Lotka–Volterra growth cycle and *s*_*t*_ is the imposed transition fraction. An analogous update was used for private-to-cheater transitions:

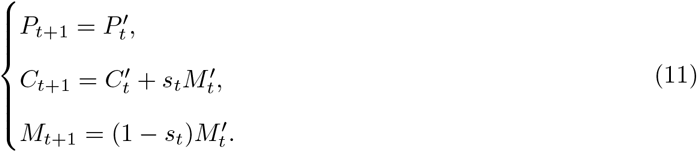

This procedure should be interpreted as a controlled phenotype-switching or strategy-perturbation simulation rather than a formal replicator-mutator model.

First, we allowed an initial shift of 30% of private metabolizers to transition into either cheater (Figure 5a) or public metabolizer (Figure 5b) strategies. Then at a rate of 10 % shift per generation, gradually. All three strategies remained coexistent and stable even after the fourth generation, and this mixed population structure persisted even when the transition rate was increased to 60%.

**Figure 5:**
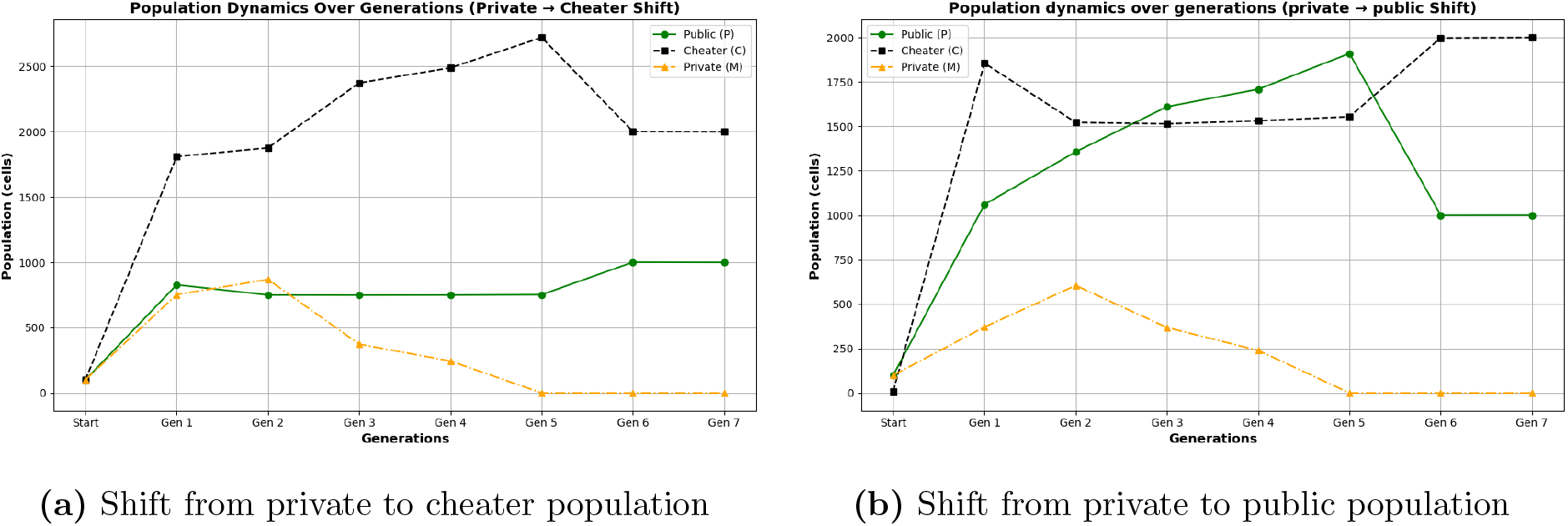
Population dynamics during evolutionary shifts from private to cheater and public strategies. (a, b) Population dynamics illustrating an initial shift of 30% of private metabolizers transitioning gradually into either cheater (a) or public metabolizer (b) strategies, with subsequent increases of 10% each generation. All three strategies remain coexistent and stable, even when up to 60% of the private metabolizer population has transitioned, demonstrating resilience and ecological stability under evolutionary changes.

Additional simulations (Figure 6) further demonstrate that when public metabolizers gain a substantial growth advantage, doubling the rate of private metabolizers, the system still supports coexistence, though at reduced equilibrium densities. This underscores the fragile ecological balance required to sustain public contributors. Although public metabolizers and cheaters may persist in small fractions, their decline coincides with an increase in total population size. These outcomes highlight the sensitivity of coexistence to fitness asymmetries and demonstrate how imbalances in growth advantages can lead to the collapse of strategic diversity within microbial communities.

**Figure 6:**
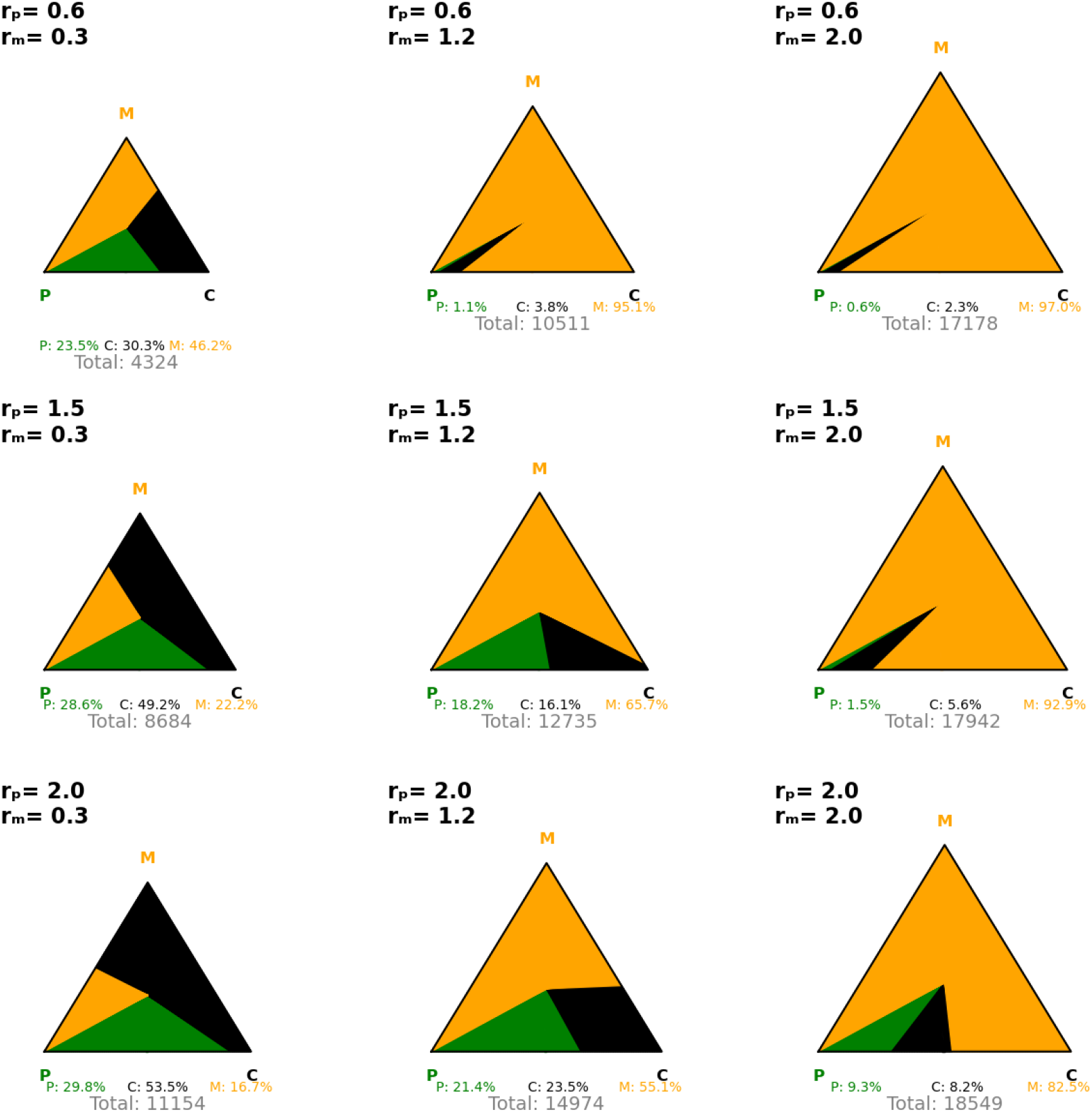
Evolutionary outcomes of a three-member three-strategy microbial community under varying payoff asymmetries in heterotrophic microbial consortia. Each triangle depicts the final population distribution of public, cheater, and private metabolizers of each generation. Colour regions indicate their relative proportions: green for *P*, black for *C*, and orange for *M*; numeric labels show final percentages and total population size. When *r*_*p*_ *> r*_*m*_, public metabolizers hold a growth advantage, cheaters exploit this benefit, and all three strategies coexist with a low total population. As *r*_*m*_ increases and surpasses *r*_*p*_, *M* dominates and total population increases, while *P* and *C* persist only in small fractions.

## 4 Discussion and Conclusion

We identified a restricted parameter regime in which public metabolizers, cheaters, and private metabolizers coexist at a positive interior equilibrium. The combined results of numerical simulations and EGT emphasize the critical conditions necessary for stable coexistence among public metabolizers, cheaters, and private metabolizers within microbial communities. Both modelling frameworks consistently reveal that maintaining a growth advantage for public metabolizers is essential for long-term coexistence. When growth rates are balanced and the cost of competition remains lower than the cost of cooperation, our simulations indicate the system tends to stabilize and steady state around a point that includes all three strategies.

However, imbalances in growth rates significantly alter community dynamics, as highlighted by numerical analyses of various initial population ratios and growth rate parameters (Figure 2,Figure S1). A higher initial ratio of cheaters consistently showed detrimental impacts on public metabolizers, suppressing their maximum population densities and long-term survival. In contrast, private metabolizers benefited indirectly under these conditions, achieving higher equilibrium densities because of decreased competition from the declining public population. These findings align with evolutionary game-theoretic predictions, supporting the conclusion that cooperative strategies are especially vulnerable to exploitation unless protected by stabilizing conditions.

Further insights emerged from evolutionary game simulations that examined population shift strategies and provided valuable insights into resilience and adaptability within microbial communities. Even when a large fraction of private metabolizers transitioned to cheating or public strategies, the system often retained all three populations, indicating a robust tolerance to evolutionary changes. Notably, even with a large fraction of private metabolizers mutating toward other strategies, it remains possible for all three strategies to persist within the community during the evolutionary phases. However, substantial shifts toward cheaters imposed significant pressure on public metabolizers, often leading to their elimination unless cooperation costs remained sufficiently low and ecological competition was constrained.

An extensive exploration of parameter spaces, including the effects of varying *α* : *β* and *β* : *α* ratios, reinforced these findings. Specifically, higher *α* : *β* ratios, representing elevated exploitation by cheaters relative to competition costs, consistently reduced the stability of cooperative interactions (Figure 3). Conversely, optimizing the balance between these parameters promoted conditions favourable for sustained coexistence. These findings highlight how the intensity and direction of competitive and exploitative pressures shape the long-term evolutionary landscape of microbial communities. Moreover, numerically, the most robust coexistence with balanced community fractions is observed when *r*_*p*_/*r*_*m*_ lies between 1.2 and 2 (Figure 6), but analytically the feasibility window extends to *r*_*p*_/*r*_*m*_ *<* 5 with the default interaction parameters (Appendix D). Analytically, the transitions between coexistence and single-strategy dominance at the boundaries of the feasibility window correspond to transcritical bifurcations [41], in which the interior equilibrium collides with and exchanges stability with a boundary equilibrium. This narrow window defines the optimal condition for maximizing overall community density while preserving diversity. If the public growth rate is double that of the private rate, the mixed community achieves maximum density. However, extreme imbalances, where one strategy significantly outpaces the others, lead to competitive exclusion. Elevated public growth results in the dominance of public metabolizers and cheaters, while significantly higher private growth leads to the extinction of the other two populations (Figure 2, Figure 6). Taken together, these results show that privatization does not necessarily lead to collapse; its effect depends on whether the private strategy lies inside or outside the coexistence window defined by growth asymmetry, exploitation strength, and density-dependent self-limitation.

The present model also extends earlier yeast sucrose and exoenzyme public-goods models. After Greig and Travisano [50] established sucrose metabolism as a canonical microbial cooperation system, experimental and theoretical studies showed that invertase-producing yeast can generate public-good dynamics in which non-producing or low-producing strains exploit glucose released by cooperators [14, 15]. Our formulation builds on this tradition but adds a third metabolic strategy, the private metabolizer, which avoids public-good sharing by internalizing resource processing. This connects our model directly to Lindsay *et al*.[32], who showed that privatization of public goods can invade cooperative systems and cause population decline. In contrast to their resource-explicit formulation, we use a reduced generalized Lotka–Volterra model to derive closed-form coexistence conditions, boundary equilibria, and stability criteria for the full public–cheater–private strategy space. The loss of resource-level mechanistic detail is therefore compensated by analytical transparency: the model identifies which combinations of measurable growth, exploitation, competition, and self-limitation parameters permit coexistence.

While multi-strategy evolutionary public goods models have been extensively studied, including optional public-goods games with loner strategies [23, 28, 29], density-dependent public-goods models [25, 27, 30], and game-environment feedback frameworks with oscillatory dynamics [31], earlier microbial exoenzyme models have also analysed cooperation and cheating in biotechnological contexts [15].

However, these approaches typically either rely on abstract payoff structures or focus on two-strategy public-good systems rather than a three-strategy public–cheater–private metabolizer framework. In contrast, our formulation introduces a mechanistically grounded three-strategy system in which interaction coefficients arise from metabolite-mediated processes and are directly linked to density-dependent population dynamics. This allows us to explicitly connect ecological interactions with evolutionary stability, and to derive conditions for coexistence within a unified eco-evolutionary framework.

The present framework is intentionally coarse-grained. It does not resolve explicit sucrose or glucose dynamics, diffusion, or spatial structure, and therefore should not be interpreted as a full mechanistic resource model. Rather, it provides an analytically tractable effective description of how growth, exploitation, competition, and self-limitation jointly shape coexistence. Likewise, our stability results are local, and the evolutionary switching simulations are intended as robustness tests rather than as a formal replicator–mutator model. More mechanistically detailed approaches, including genome-scale metabolic models with dynamic flux-balance analysis, such as COMETS [51] or BacArena [52], can resolve strain-specific metabolism, metabolite exchange, and spatial or community-level flux constraints. Such models are powerful for predicting feasible metabolic interactions in specific consortia, but they often require extensive genome annotations, uptake parameters, and environmental constraints, and usually do not yield closed-form coexistence or stability conditions. Our reduced formulation is therefore complementary: it sacrifices intracellular metabolic detail in order to obtain analytically transparent criteria for coexistence, boundary collapse, and parameter sensitivity. In this sense, the model is intended as a conceptual and experimentally parameterizable layer between abstract public-goods games and full genome-scale community simulations.

Finally, from a biotechnological perspective, the model yields several practical design principles for creating and managing microbial communities. First, public-good producers should not merely be introduced into a consortium. Stable coexistence is contingent upon a precise interplay between growth advantages, cooperation costs, and population structure. Hence, their growth advantage must be sufficient to compensate for exploitation by cheaters, but not so high that private metabolizers are excluded. Our findings suggest that targeted interventions, such as modulating nutrient levels, introducing feedback-limited gene circuits, or engineering metabolic constraints, can reinforce this balance and sustain desired community states. By identifying parameter regimes that support robustness and diversity, this work contributes to the rational design of synthetic consortia and the ecological stabilization of natural microbial systems. These principles are directly relevant for synthetic microbial consortia in which extracellular enzymes, cross-feeding, or shared metabolic intermediates are used to distribute metabolic labour across community members. Such designs are increasingly explored in bioproduction, environmental bioremediation, and microbiome engineering, where community performance depends not only on the metabolic capacity of individual strains but also on maintaining stable interactions, functional diversity, and resistance to collapse [53, 54, 55, 56, 57]. Together, these results support the use of low-dimensional, experimentally parameterizable models as design tools for predicting when engineered microbial consortia will maintain cooperation, tolerate cheaters, or collapse into privatized metabolism.

## Acknowledgment

This work has been funded by the Deutsche Forschungsgemeinschaft (DFG, German Research Foundation) – SFB1535 - Project ID 458090666.

## Supplementary Material

**Figure S1:**
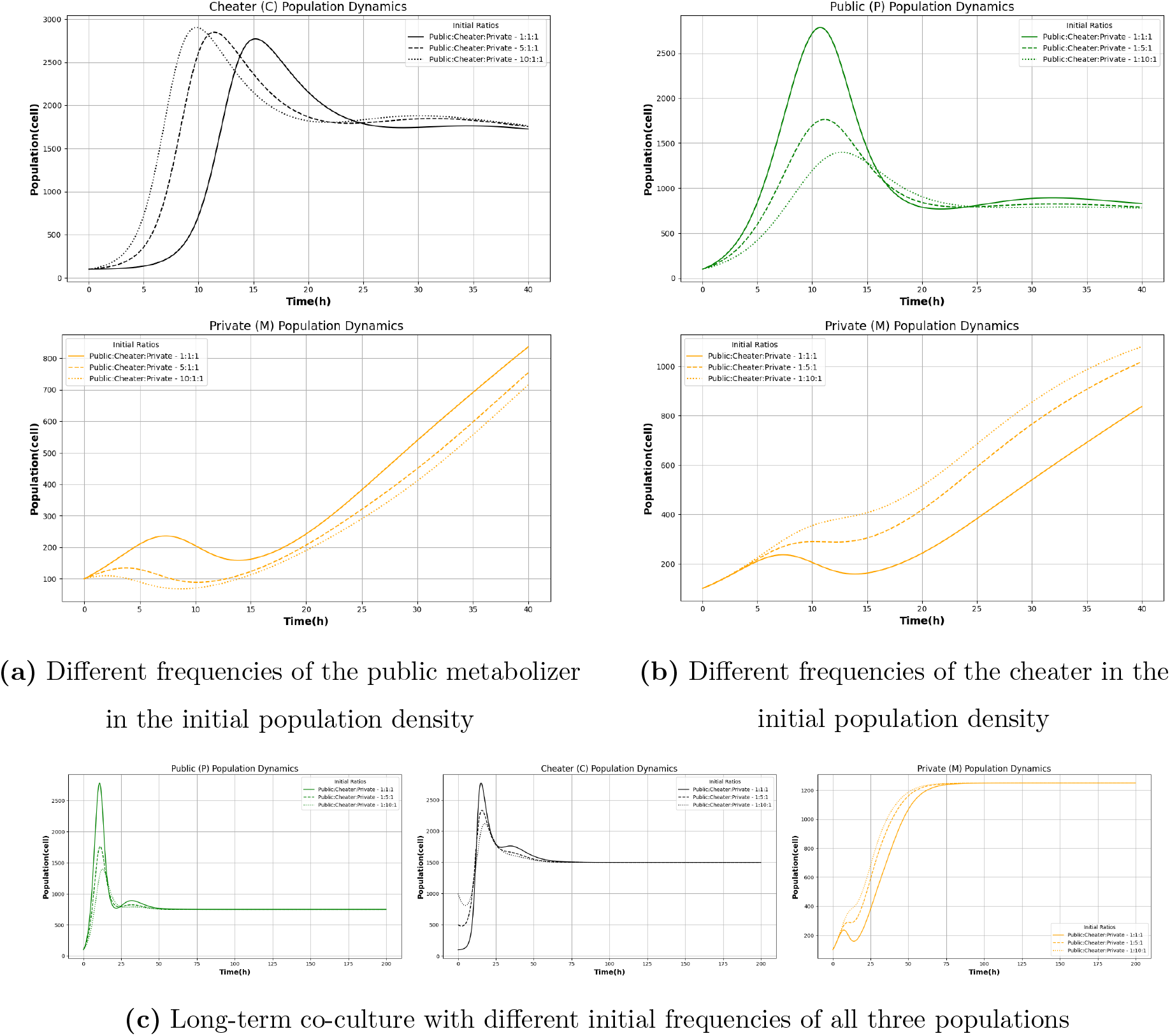
The population dynamics of all three community members depend on the initial frequencies. (a) and (b) illustrate how early dynamics differ with changing initial abundances of the public metabolizer and the cheater, respectively. (c) shows that despite these initial differences, all communities converge to a similar steady state in long-term coculture. This indicates that long-term coexistence is largely independent of the initial population composition.

### A Parameters sensitivity analysis

In microbial ecosystems, sensitivity analyses are essential to identify which biological interactions, such as cooperation or competition, most critically influence population stability and composition. The relative influence of each parameter on system behaviour can be quantified by introducing perturbations to key parameters, such as *α*, and examining the resulting changes in steady-state population densities. To assess the relative impact of a given parameter on a population (*X* ∈ {*P, C, M*}), the deviation between the perturbed and baseline population trajectories is integrated over time, as expressed by the following relation:

**Figure S2:**
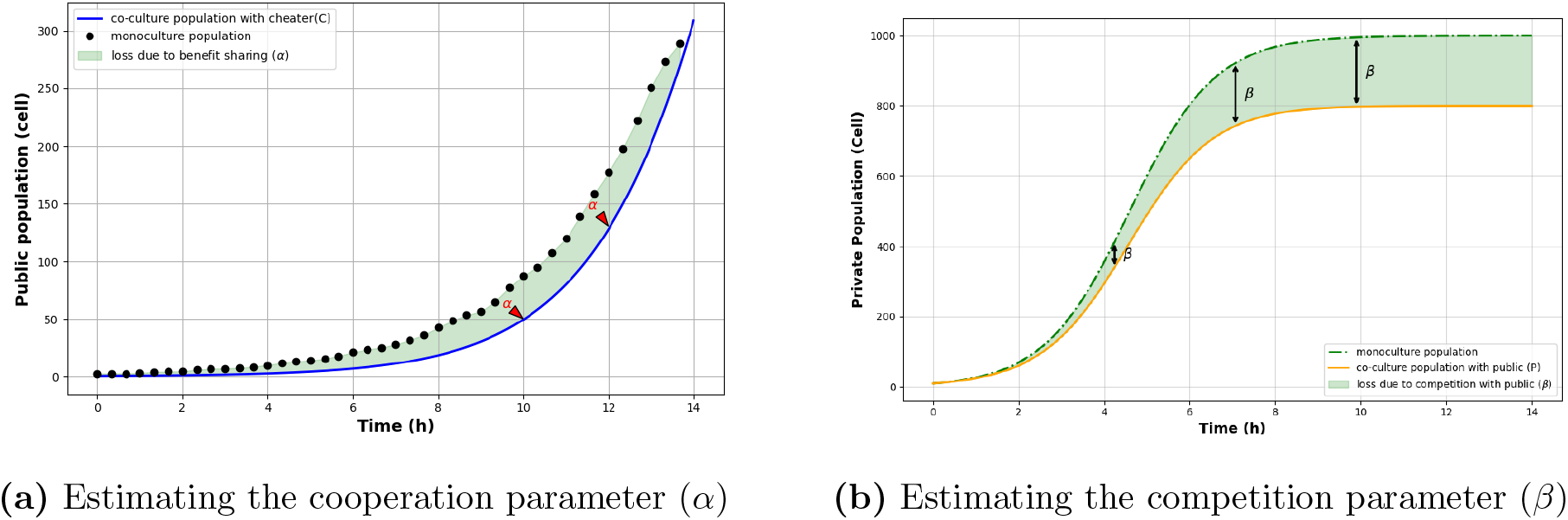
Estimating interaction parameters from mono- and co-cultures. (a) To estimate the cooperation parameter (*α*), we compare the growth trajectory of the public metabolizer (*P*) in monoculture (black points) with its growth in co-culture with the cheater (*C*) (blue line). The area between the curves (shaded green) quantifies the reduction in population size due to shared public goods, highlighting the fitness cost of exploitation. (b) To estimate the competition parameter(*β*), we compare the growth of the private metabolizer (*M*) in monoculture (green dashed line) and co-culture with the public metabolizer (*P*) (orange line). The shaded area illustrates the population loss attributable to indirect competition for sucrose resources. These pairwise measurements enable data-informed parametrization of our mathematical model.

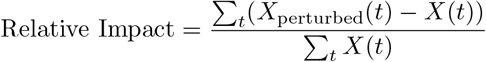

where *X*_perturbed_(*t*) denotes the population trajectory under parameter perturbation, and *X*(*t*) represents the steadystate baseline trajectory. This formulation captures the cumulative deviation caused by the perturbation, offering a time-integrated measure of sensitivity over the simulation period. As parameter sensitivity may vary depending on the magnitude of perturbation, the average impact is computed across multiple perturbation steps:

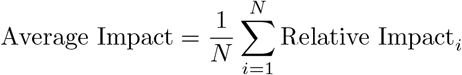

Here, *N* is the total number of perturbation steps. Averaging across these steps helps reduce the influence of noise or isolated fluctuations and provides a more robust estimate of each parameter’s influence. This approach provides a robust measure of a parameter’s influence, minimising the effects of noise or extreme variations. To ensure consistency and comparability across different parameters and populations, the computed impacts are normalised to a standard scale within the range [−1, 1].

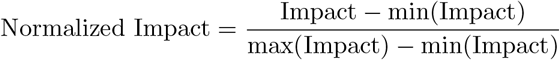

Normalisation allows for direct comparison between parameters, ensuring that those with stronger relative effects are identified, even when absolute magnitudes differ due to different units or scales.

**Figure S3:**
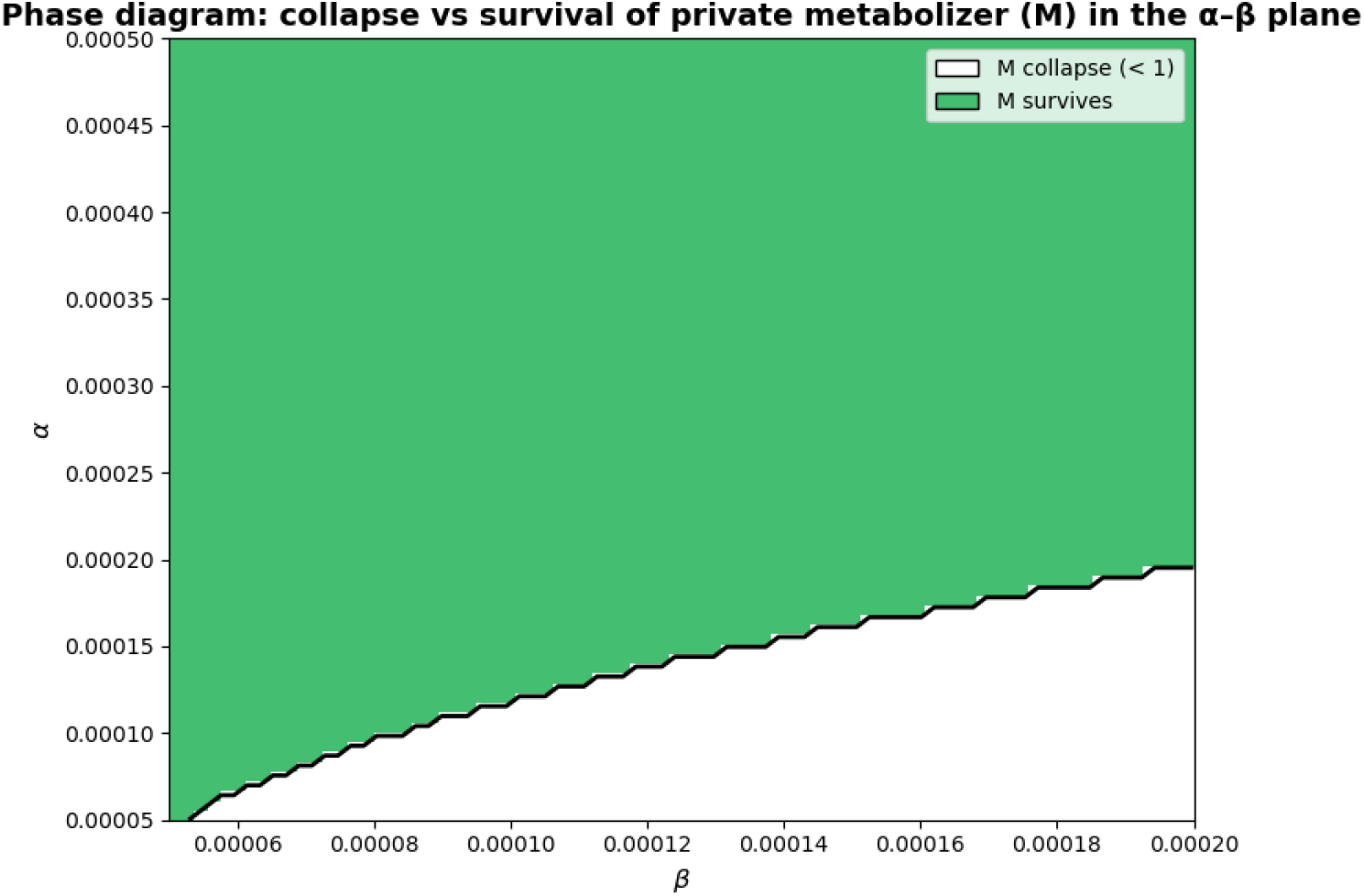
Phase diagram: collapse vs survival of private metabolizer (*M*) in the *α*-*β* plane. The parameters are varied in a range between 0.5 and 5 times baseline. Initial starting conditions were set to 100 for all three players.

**Figure S4:**
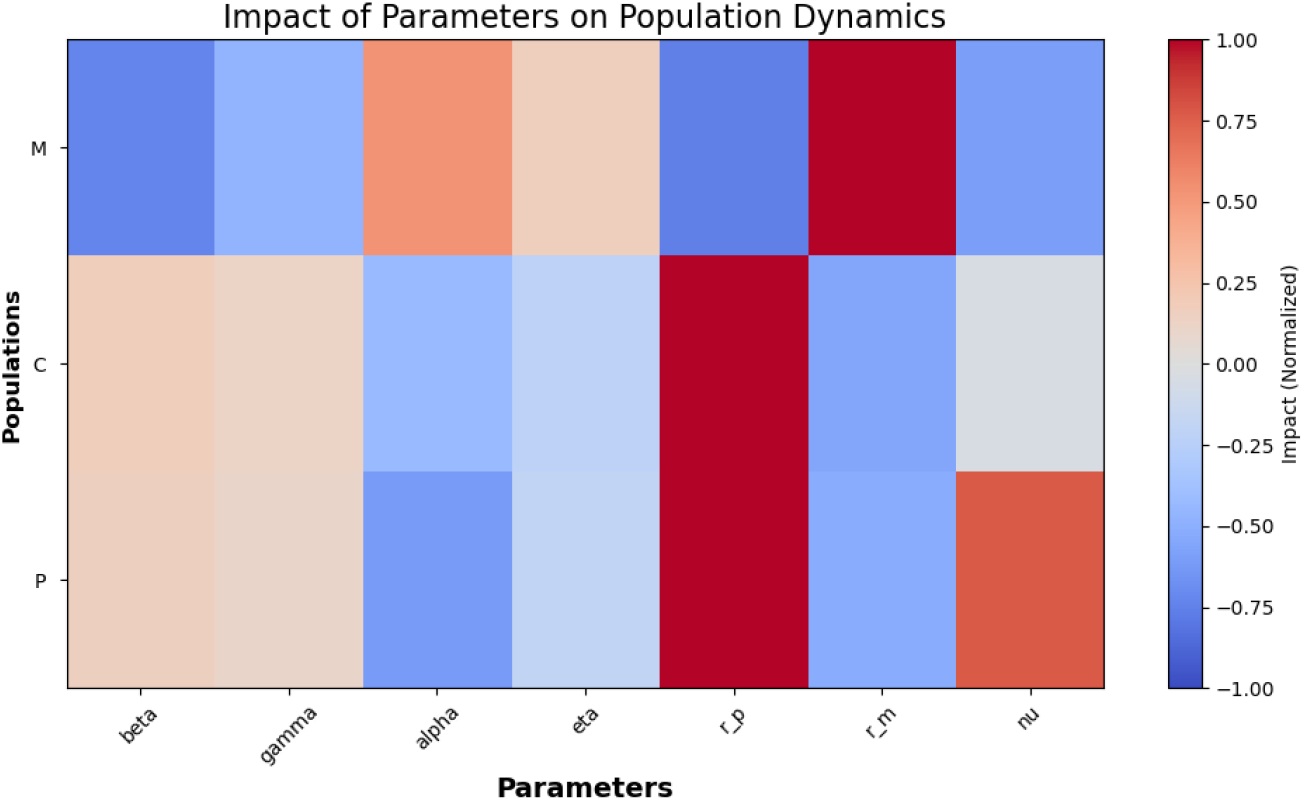
Parameter sensitivity of population dynamics assessed via perturbation analysis. Each model parameter was independently scaled over a predefined range (1–3 x baseline), and the resulting trajectories were simulated from fixed initial conditions. The heatmap shows the mean relative change in cumulative population abundance (time-integrated density) for public metabolizers (*P*), cheaters (*C*), and private metabolizers (*M*), normalized to the unperturbed baseline simulation. Positive values indicate an increase in total population abundance under parameter perturbation, while negative values indicate a decrease. The analysis provides a coarse-grained approximation of parameter influence analogous to metabolic control analysis, highlighting differential sensitivity of each population to model parameters.

### B One-at-a-time parameter sensitivity

Relative changes in steady-state abundances (*P*^*^, *C*^*^, *M*^*^) when each parameter is varied individually by the indicated factor while all other parameters remain at their default values (Table 1). Only parameter combinations yielding a positive interior equilibrium (*D >* 0 and *P* ^*^, *C*^*^, *M*^*^ *>* 0) are reported.

**Table S1:**
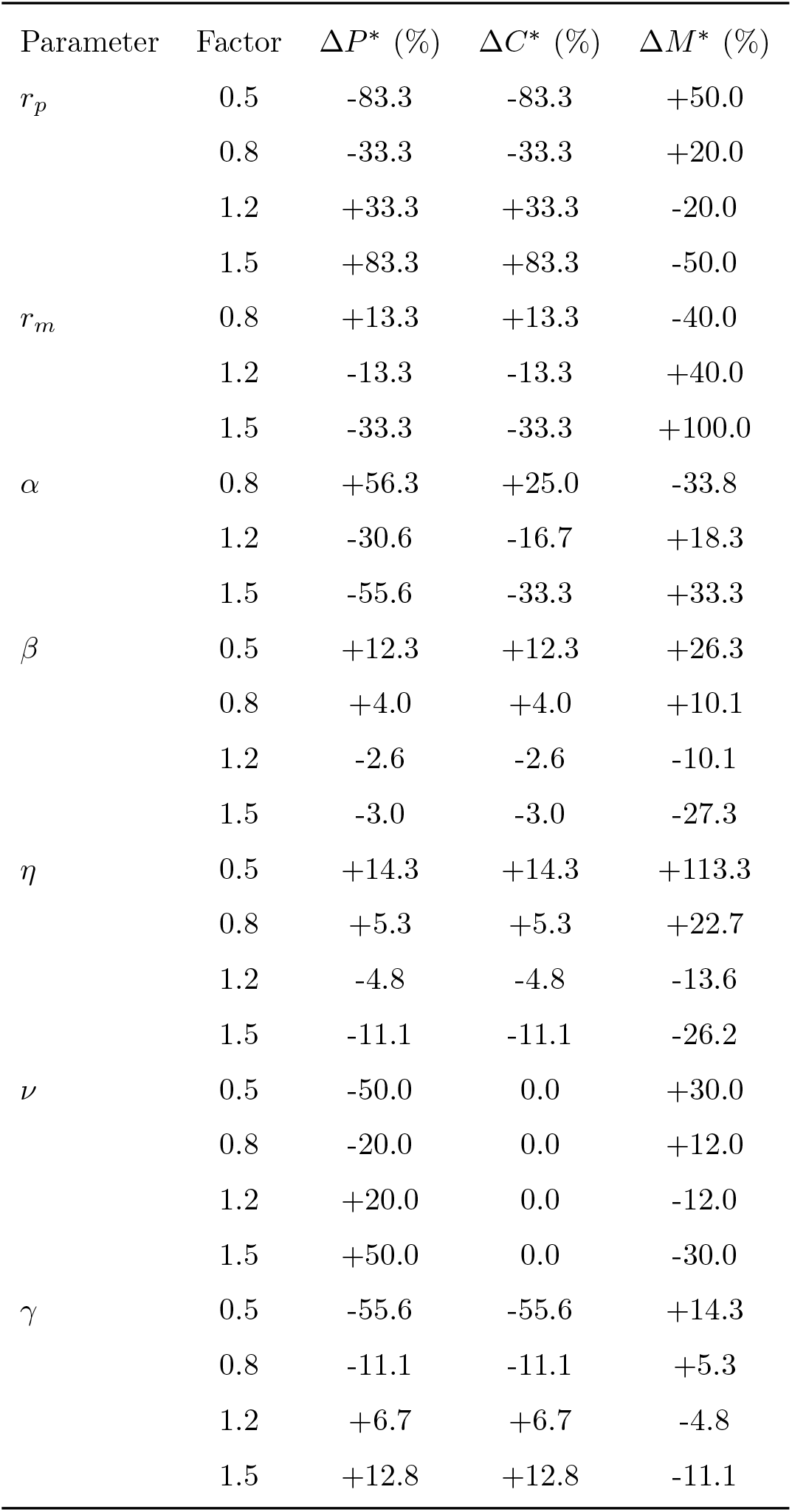
Relative change in equilibrium abundances under parameter perturbations.

| Parameter | Factor | $\Delta P^*$ (%) | $\Delta C^*$ (%) | $\Delta M^*$ (%) |
| --- | --- | --- | --- | --- |
| $r_p$ | 0.5 | -83.3 | -83.3 | +50.0 |
|  | 0.8 | -33.3 | -33.3 | +20.0 |
|  | 1.2 | +33.3 | +33.3 | -20.0 |
|  | 1.5 | +83.3 | +83.3 | -50.0 |
| $r_m$ | 0.8 | +13.3 | +13.3 | -40.0 |
|  | 1.2 | -13.3 | -13.3 | +40.0 |
|  | 1.5 | -33.3 | -33.3 | +100.0 |
| $\alpha$ | 0.8 | +56.3 | +25.0 | -33.8 |
|  | 1.2 | -30.6 | -16.7 | +18.3 |
|  | 1.5 | -55.6 | -33.3 | +33.3 |
| $\beta$ | 0.5 | +12.3 | +12.3 | +26.3 |
|  | 0.8 | +4.0 | +4.0 | +10.1 |
|  | 1.2 | -2.6 | -2.6 | -10.1 |
|  | 1.5 | -3.0 | -3.0 | -27.3 |
| $\eta$ | 0.5 | +14.3 | +14.3 | +113.3 |
|  | 0.8 | +5.3 | +5.3 | +22.7 |
|  | 1.2 | -4.8 | -4.8 | -13.6 |
|  | 1.5 | -11.1 | -11.1 | -26.2 |
| $\nu$ | 0.5 | -50.0 | 0.0 | +30.0 |
|  | 0.8 | -20.0 | 0.0 | +12.0 |
|  | 1.2 | +20.0 | 0.0 | -12.0 |
|  | 1.5 | +50.0 | 0.0 | -30.0 |
| $\gamma$ | 0.5 | -55.6 | -55.6 | +14.3 |
|  | 0.8 | -11.1 | -11.1 | +5.3 |
|  | 1.2 | +6.7 | +6.7 | -4.8 |
|  | 1.5 | +12.8 | +12.8 | -11.1 |

### C Interior evolutionary equilibrium: derivation, uniqueness, and payoff equivalence

Setting the original ODE system (Eq. 1) to 0:

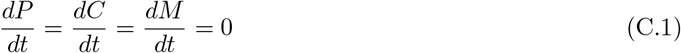

and requiring

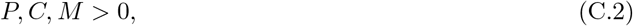

we obtain the following linear system after dividing each equation by the respective positive population density:

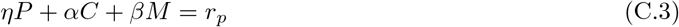

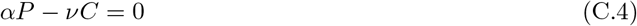

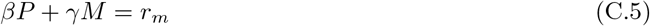

From Eq. (C.4),

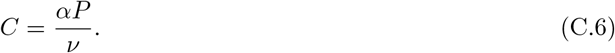

From Eq. (C.5),

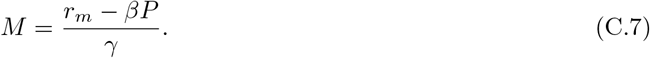

Substituting Eqs. (C.6) and (C.7) into Eq. (C.3) and rearranging gives:

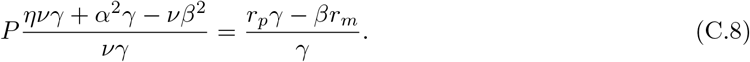

Since

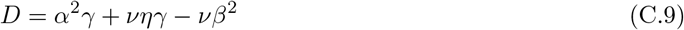

the unique solution is:

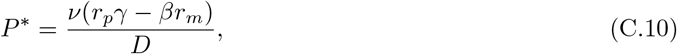

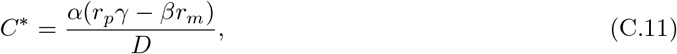

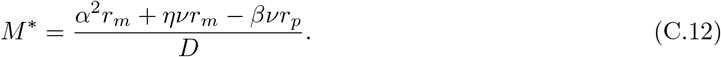

Because the three steady-state conditions reduce to a system of three linear equations in *P, C, M*, this solution is unique whenever *D*≠ 0, precluding a second interior equilibrium.

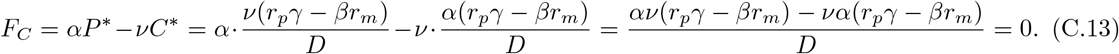

For *F*_*M*_, using *M*^*^ = (*r*_*m*_ − *βP*^*^)*/γ* from Eq. (C.5):

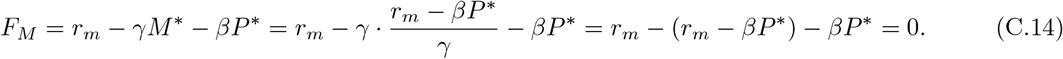

For *F*_*P*_, substituting *P* ^*^, *C*^*^, *M*^*^ and multiplying through by *D*:

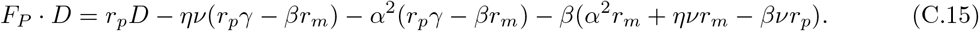

Expanding and collecting terms in *r*_*p*_ and *r*_*m*_, all coefficients cancel exactly, giving *F*_*P*_ · *D* = 0, and hence *F*_*P*_ = 0.

Thus *F*_*P*_ = *F*_*C*_ = *F*_*M*_ = 0 holds identically at (*P*^*^, *C*^*^, *M*^*^).

### D Proof of local asymptotic stability

The full Jacobian of the Lotka–Volterra system is

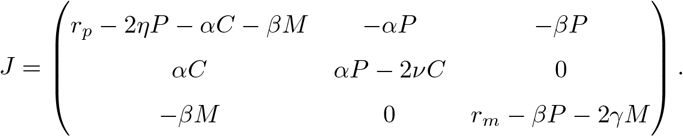

At the interior equilibrium (*P*^*^, *C*^*^, *M*^*^), this simplifies to the evaluated Jacobian

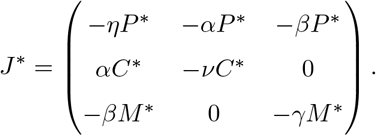

The trace is strictly negative,

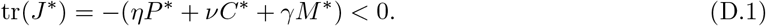

The characteristic polynomial of *J*^*^ is

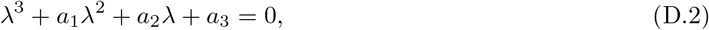

with coefficients

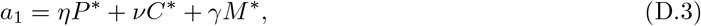

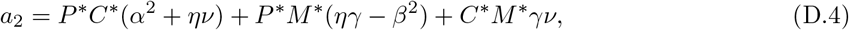

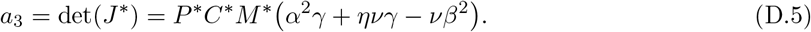

Defining

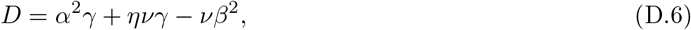

one has *a*_3_ = *P*^*^*C*^*^*M*^*^*D*, so *a*_3_ *>* 0 whenever *P*^*^, *C*^*^, *M*^*^ *>* 0 and *D >* 0.

By the Routh–Hurwitz criterion for cubic polynomials, local asymptotic stability is ensured if

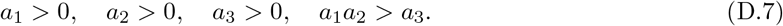

The first and third conditions follow directly from positivity of the equilibrium densities and *D >* 0. The coefficient *a*_2_ is positive provided

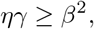

which ensures that all pairwise interaction contributions are stabilizing. Under this condition, all coefficients are strictly positive, and the remaining Routh–Hurwitz inequality *a*_1_*a*_2_ *> a*_3_ is satisfied.

Therefore, the interior equilibrium (*P* ^*^, *C*^*^, *M*^*^) is locally asymptotically stable provided it exists with positive components and the interaction parameters satisfy *D* = *α*^2^*γ* +*νηγ* −*νβ*^2^ *>* 0 (which holds whenever *ηγ > β*^2^, a condition satisfied by the default parameters and explored ranges). The requirements *P*^*^ *>* 0 and *C*^*^ *>* 0 yield the lower bound

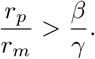

The requirement *M*^*^ *>* 0 yields the upper bound

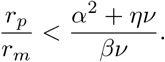

Thus, stable tripartite coexistence is possible only inside the window

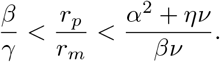

With the default parameter values (*α* = 0.0002, *β* = 0.0001, *γ* = 0.0001, *η* = 0.0001,*ν* = 0.0001), this window evaluates to approximately 1 *< r*_*p*_*/r*_*m*_ *<* 5.

### E Boundary equilibria

To fully characterize the dynamics, we enumerate all equilibria of system (1). In addition to the interior fixed point given in Eq. (8), the system admits the following boundary equilibria.

The trivial equilibrium is

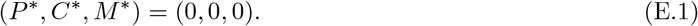

It always exists and is unstable whenever *r*_*p*_ *>* 0 or *r*_*m*_ *>* 0.

The *M*-only equilibrium is

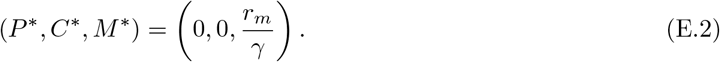

Here, *P* and *C* are absent, while *M* reaches its carrying capacity. This equilibrium exists for all positive parameter values.

The *P*-only equilibrium is

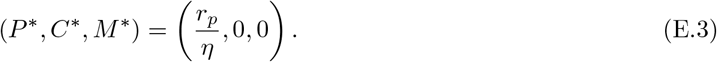

Here, only the public metabolizer persists. This equilibrium exists for all positive parameter values.

The *P*–*C* subsystem equilibrium is

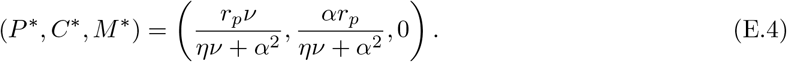

At this equilibrium, the public metabolizer and cheater coexist in the absence of the private metabolizer.

The *P*–*M* subsystem equilibrium exists when

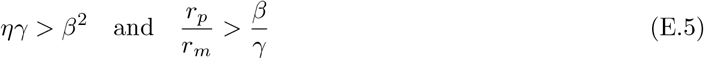

and therefore is equal to:

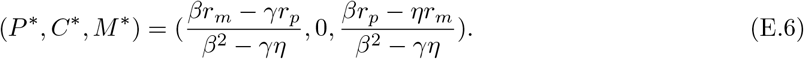

There is no *C*-only or *C*–*M* equilibrium. Since

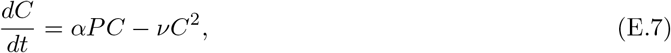

setting *P* = 0 gives

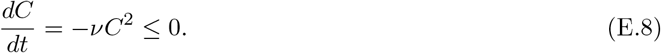

Thus, cheaters cannot persist without public metabolizers.

## References

[1] Hillman ET, Lu H, Yao T, Nakatsu CH. 2017 Microbial ecology along the gastrointestinal tract. Microbes and environments 32, 300–313.

[2] Amato P, Joly M, Besaury L, Oudart A, Taib N, Moné AI, Deguillaume L, Delort AM, Debroas D. 2017 Active microorganisms thrive among extremely diverse communities in cloud water. PloS one 12, e0182869.

[3] Solbrig OT. 1991 The origin and function of biodiversity. Environment: Science and Policy for Sustainable Development 33, 16–38.

[4] Rutigliano FA, D’Ascoli R, De Santo AV. 2004 Soil microbial metabolism and nutrient status in a Mediterranean area as affected by plant cover. Soil Biology and Biochemistry 36, 1719–1729.

[5] Matuszyńska A, Ebenhöh O, Zurbriggen MD, Ducat DC, Axmann IM. 2024 A new era of synthetic biology—microbial community design. Synthetic Biology 9, ysae011. (10.1093/synbio/ysae011)

[6] Mkilima T. 2025 Engineering artificial microbial consortia for personalized gut microbiome modulation and disease treatment. Annals of the New York Academy of Sciences.

[7] Rafeeq H, Afsheen N, Rafique S, Arshad A, Intisar M, Hussain A, Bilal M, Iqbal HM. 2023 Genetically engineered microorganisms for environmental remediation. Chemosphere 310, 136751.

[8] Ruan Z, Chen K, Cao W, Meng L, Yang B, Xu M, Xing Y, Li P, Freilich S, Chen C et al. 2024 Engineering natural microbiomes toward enhanced bioremediation by microbiome modeling. Nature Communications 15, 4694.

[9] Jimenez P, Scheuring I. 2021 Density-dependent private benefit leads to bacterial mutualism. Evolution 75, 1619–1635. (10.1111/evo.14241)

[10] Srinivasan S, Jnana A, Murali TS. 2024 Modeling Microbial Community Networks: Methods and Tools for Studying Microbial Interactions. Microbial Ecology 87, 56. (10.1007/s00248-024-02370-7)

[11] Czárán TL, Hoekstra RF. 2009 Microbial Communication, Cooperation and Cheating: Quorum Sensing Drives the Evolution of Cooperation in Bacteria. PLoS ONE 4, e6655. (10.1371/journal.pone.0006655)

[12] Czárán TL et al. 2024 Cue-driven microbial cooperation and communication: evolving quorum sensing with honest signaling. BMC Biology 22, 73. (10.1186/s12915-024-01855-0)

[13] Scherlach K, Hertweck C. 2018 Mediators of mutualistic microbe–microbe interactions. Natural Product Reports 35, 303–308.

[14] Gore J, Youk H, Van Oudenaarden A. 2009 Snowdrift game dynamics and facultative cheating in yeast. Nature 459, 253–256. (10.1038/nature07921)

[15] Schuster S, Kreft JU, Brenner N, Wessely F, Theißen G, Ruppin E, Schroeter A. 2010 Cooperation and cheating in microbial exoenzyme production - Theoretical analysis for biotechnological applications. Biotechnology Journal 5, 751–758. (10.1002/biot.200900303)

[16] Mostafa F, Kruger A, Nies T, Frunzke J, Schipper K, Matuszyńska A. 2024 Microbial markets: socio-economic perspective in studying microbial communities. microLife p. uqae016.

[17] Rainey PB, Rainey K. 2003 Evolution of cooperation and conflict in experimental bacterial populations. Nature 425, 72–74.

[18] West SA, Griffin AS, Gardner A, Diggle SP. 2006 Social evolution theory for microorganisms. Nature reviews microbiology 4, 597–607.

[19] Frey E. 2010 Evolutionary game theory: Theoretical concepts and applications to microbial communities. Physica A: Statistical Mechanics and its Applications 389, 4265–4298.

[20] Garde R, Ewald J, Kovács ÁT, Schuster S. 2020 Modelling population dynamics in a unicellular social organism community using a minimal model and evolutionary game theory. Open Biology 10, 200206.

[21] Leimar O, McNamara JM. 2023 Game theory in biology: 50 years and onwards. Philosophical Transactions of the Royal Society B 378, 20210509.

[22] Grodwohl JB, Parker GA. 2023 The early rise and spread of evolutionary game theory: perspectives based on recollections of early workers. Philosophical Transactions of the Royal Society B 378, 20210493.

[23] Hauert C, De Monte S, Hofbauer J, Sigmund K. 2002 Replicator dynamics for optional public good games. Journal of Theoretical Biology 218, 187–194. (10.1006/jtbi.2002.3067)

[24] Hauert C, Szabó G. 2003 Prisoner’s Dilemma and Public Goods Games in Different Geometries: Compulsory versus voluntary interactions. Complexity 8, 31–38. (10.1002/cplx.10084)

[25] Hauert C. 2008 Ecological public goods games: cooperation and bifurcation. Theoretical Population Biology 73, 257–263. (10.1016/j.tpb.2007.11.004)

[26] Smith P, Schuster M. 2019 Public goods and cheating in microbes. Current biology 29, R442–R447.

[27] Requejo RJ, Camacho J. 2012 Coexistence of Cooperators and Defectors in Well Mixed Populations Mediated by Resource Limitation. Physical Review Letters 108, 038102. (10.1103/Phys-RevLett.108.038102)

[28] Hauert C, De Monte S, Hofbauer J, Sigmund K. 2002 Volunteering as Red Queen Mechanism for Cooperation in Public Goods Games. Science 296, 1129–1132. (10.1126/science.1070582)

[29] Doebeli M, Hauert C, Timothy K. 2004 The evolutionary origin of cooperators and defectors. Science 306, 859–862. (10.1126/science.1101456)

[30] Archetti M, Scheuring I. 2011 Coexistence of cooperation and defection in public goods games. Evolution 65, 1140–1148. (10.1111/j.1558-5646.2010.01185.x)

[31] Weitz JS, Eksin C, Paarporn K, Brown SP, Ratcliff WC. 2016 An oscillating tragedy of the commons in replicator dynamics with game-environment feedback. Proceedings of the National Academy of Sciences 113. (10.1073/pnas.1604096113)

[32] Lindsay RJ, Pawlowska BJ, Gudelj I. 2019 Privatization of public goods can cause population decline. Nature Ecology & Evolution 3, 1206–1216. (10.1038/s41559-019-0944-9)

[33] Germerodt S, Bohl K, Lück A, Pande S, Schröter A, Kaleta C, Schuster S, Kost C. 2016 Pervasive selection for cooperative cross-feeding in bacterial communities. PLoS computational biology 12, e1004986.

[34] Xenophontos C, Harpole WS, Küsel K, Clark AT. 2022 Cheating promotes coexistence in a two-species one-substrate culture model. Frontiers in Ecology and Evolution 9, 786006.

[35] Rankin DJ, Bargum K, Kokko H. 2007 The tragedy of the commons in evolutionary biology. Trends in ecology & evolution 22, 643–651.

[36] Liu M, Wild G, West SA. 2023 Equilibria and oscillations in cheat–cooperator dynamics. Evolution Letters 7, 339–350.

[37] Schuster M, Foxall E, Finch D, Smith H, De Leenheer P. 2017 Tragedy of the commons in the chemostat. PLOS ONE 12, e0186119. (10.1371/journal.pone.0186119)

[38] Lotka AJ. 1920 Analytical note on certain rhythmic relations in organic systems. Proceedings of the National Academy of Sciences 6, 410–415.

[39] Volterra V. 1926 Variazioni e fluttuazioni del numero d’individui in specie animali conviventi. Società anonima tipografica “Leonardo da Vinci”.

[40] Murray JD. 2007 Mathematical biology: I. An introduction vol. 17. Springer Science & Business Media.

[41] Strogatz SH. 2018 Nonlinear dynamics and chaos: with applications to physics, biology, chemistry, and engineering. CRC press.

[42] Neumann G, Schuster S. 2007 Continuous model for the rock–scissors–paper game between bacteriocin producing bacteria. Journal of Mathematical Biology 54, 815–846. (10.1007/s00285-006-0065-3)

[43] Křivan V, Galanthay TE, Cressman R. 2018 Beyond replicator dynamics: From frequency to density dependent models of evolutionary games. Journal of Theoretical Biology 455, 232–248. (10.1016/j.jtbi.2018.07.003)

[44] Smith JM, Price GR. 1973 The logic of animal conflict. Nature 246, 15–18.

[45] Remien CH, Eckwright MJ, Ridenhour BJ. 2021 Structural identifiability of the generalized Lotka–Volterra model for microbiome studies. Royal Society Open Science 8, 201378. (10.1098/rsos.201378)

[46] Uppal G, Vural DC. 2024 On the possibility of engineering social evolution in microfluidic environments. Biophysical Journal 123, 407–419.

[47] Koschwanez JH, Foster KR, Murray AW. 2011 Sucrose Utilization in Budding Yeast as a Model for the Origin of Undifferentiated Multicellularity. PLOS Biology 9, 1–10. (10.1371/journal.pbio.1001122)

[48] van Aalst M, Lahlou A, Hassan T, Gaultier W, Colliaux D, Matuszyńska A. 2026 Web-based collaborative model development in interdisciplinary consortia. PLOS Biology. Accepted for publication.

[49] Wang Q, He N, Chen X. 2018 Replicator dynamics for public goods game with resource allocation in large populations. Applied Mathematics and Computation 328, 162–170.

[50] Greig D, Travisano M The Prisoner’s Dilemma and polymorphism in yeast SUC genes. 271. (10.1098/rsbl.2003.0083)

[51] Dukovski I, Bajić D, Chacón JM, Quintin M, Vila JCC, Sulheim S, Pacheco AR, Bernstein DB, Riehl WJ, Korolev KS, Sanchez A, Harcombe WR, Segrè D A metabolic modeling platform for the computation of microbial ecosystems in time and space (COMETS). 16, 5030–5082. (10.1038/s41596-021-00593-3)

[52] Bauer E, Zimmermann J, Baldini F, Thiele I, Kaleta C. 2017 BacArena: Individual-based metabolic modeling of heterogeneous microbes in complex communities. PLOS Computational Biology 13, 1–22. (10.1371/journal.pcbi.1005544)

[53] McCarty NS, Ledesma-Amaro R Synthetic Biology Tools to Engineer Microbial Communities for Biotechnology. 37, 181–197. (10.1016/j.tibtech.2018.11.002)

[54] Roell GW, Zha J, Carr RR, Koffas MA, Fong SS, Tang YJ Engineering microbial consortia by division of labor. 18, 35. (10.1186/s12934-019-1083-3)

[55] Li X, Zhou Z, Li W, Yan Y, Shen X, Wang J, Sun X, Yuan Q Design of stable and self-regulated microbial consortia for chemical synthesis. 13, 1554. (10.1038/s41467-022-29215-6)

[56] van Leeuwen PT, Brul S, Zhang J, Wortel MT Synthetic microbial communities (SynComs) of the human gut: design, assembly, and applications. 47, fuad012. (10.1093/femsre/fuad012)

[57] Ruan Z, Chen K, Cao W, Meng L, Yang B, Xu M, Xing Y, Li P, Freilich S, Chen C, Gao Y, Jiang J, Xu X Engineering natural microbiomes toward enhanced bioremediation by microbiome modeling. 15, 4694. (10.1038/s41467-024-49098-z)

